# Morpho-electric and transcriptomic divergence of the layer 1 interneuron repertoire in human versus mouse neocortex

**DOI:** 10.1101/2022.10.24.511199

**Authors:** Thomas Chartrand, Rachel Dalley, Jennie Close, Natalia A. Goriounova, Brian R. Lee, Rusty Mann, Jeremy A. Miller, Gabor Molnar, Alice Mukora, Lauren Alfiler, Katherine Baker, Trygve E. Bakken, Jim Berg, Darren Bertagnolli, Thomas Braun, Krissy Brouner, Tamara Casper, Eva Adrienn Csajbok, Nick Dee, Tom Egdorf, Rachel Enstrom, Anna A. Galakhova, Amanda Gary, Emily Gelfand, Jeff Goldy, Kristen Hadley, Tim S. Heistek, DiJon Hill, Nik Jorstad, Lisa Kim, Agnes Katalin Kocsis, Lauren Kruse, Michael Kunst, Gabriela Leon, Brian Long, Matthew Mallory, Medea McGraw, Delissa McMillen, Erica J. Melief, Norbert Mihut, Lindsay Ng, Julie Nyhus, Victoria Omstead, Zoltan Peterfi, Alice Pom, Lydia Potekhina, Ramkumar Rajanbabu, Marton Rozsa, Augustin Ruiz, Joanna Sandle, Susan M. Sunkin, Ildiko Szots, Michael Tieu, Martin Toth, Jessica Trinh, Sara Vargas, David Vumbaco, Grace Williams, Julia Wilson, Zizhen Yao, Pal Barzo, Charles Cobbs, Richard G. Ellenbogen, Luke Esposito, Manuel Ferreira, Nathan W. Gouwens, Benjamin Grannan, Ryder P. Gwinn, Jason S. Hauptman, Tim Jarsky, C.Dirk Keene, Andrew L. Ko, Christof Koch, Jeffrey G. Ojemann, Anoop Patel, Jacob Ruzevick, Daniel L. Silberberg, Kimberly Smith, Staci A. Sorensen, Bosiljka Tasic, Jonathan T. Ting, Jack Waters, Christiaan P.J. de Kock, Huib D. Mansvelder, Gabor Tamas, Hongkui Zeng, Brian Kalmbach, Ed S. Lein

## Abstract

Neocortical layer 1 (L1) is a site of convergence between pyramidal neuron dendrites and feedback axons where local inhibitory signaling can profoundly shape cortical processing. Evolutionary expansion of human neocortex is marked by distinctive pyramidal neuron types with extensive branching in L1, but whether L1 interneurons are similarly diverse is underexplored. Using patch-seq recordings from human neurosurgically resected tissues, we identified four transcriptomically defined subclasses, unique subtypes within those subclasses and additional types with no mouse L1 homologue. Compared with mouse, human subclasses were more strongly distinct from each other across all modalities. Accompanied by higher neuron density and more variable cell sizes compared with mouse, these findings suggest L1 is an evolutionary hotspot, reflecting the increasing demands of regulating the expanding human neocortical circuit.

**One Sentence Summary:** Using transcriptomics and morpho-electric analyses, we describe innovations in human neocortical layer 1 interneurons.

## Main Text

Neocortical layer 1 (L1) is implicated in several higher order brain functions, including state modulation (*1*), learning (*2–5*), sensory perception (*6*), and consciousness (*7*). The neural circuitry that mediates these functions consists of converging pyramidal cell dendrites, long-range axons originating from thalamic, cortical and neuromodulatory regions and axons from local GABAergic interneurons (*8*). Much of this inhibitory input arises from neurons with cell bodies in L1, an entirely GABAergic cell population with distinct developmental origins (*9, 10*). Emerging evidence suggests that these L1 interneurons profoundly shape cortical processing and that diversity within this population is linked to diversity of function (*11, 12*). As such, the L1 interneuron repertoire is a potential site of evolutionary divergence that could contribute to specialized cortical function in humans and other primates. In rodents, a progression of classification schemes for L1 neurons (*13–18*) has evolved towards a view of 4 canonical types based on molecular markers (*19*), but the robustness of this scheme, both across modalities and across species, remains unclear (particularly in human and non-human primates). Indeed, the observation of a ‘rosehip’ cell type found in human and not mouse neocortex (*20*) highlights the importance of studying human L1 to identify potential species specializations and to relate mouse literature to human L1 cell types and function.

Traditionally, L1 cell types have been defined by their morphology, sublaminar location, intrinsic membrane properties, and a handful of marker genes. However, applying distinctive features from rodent to define and study human cell types can be tenuous. Single-cell whole transcriptome data, on the other hand, can be leveraged to define cross-species cell type homologies (*21*, *22*) and reveal genetic and phenotypic diversity obscured by the marker gene approach (*23, 24*), as observed *in vivo* in mouse L1 (*11*). The patch-seq technique (*25, 26*), combining patch-clamp electrophysiology, nuclear RNA sequencing, and morphological reconstruction from the same neuron, gives us unprecedented ability to reveal cell type diversity in human L1. We leverage this multimodal data to provide new perspective on cell-type distinctions previously proposed from a subset of modalities, make principled cross-species comparisons, and robustly identify distinct phenotypes found in human L1 across modalities.

## Results

### L1 patch-seq pipeline and transcriptomic references

To structure analysis of L1 cell types, we used transcriptomic types (t-types) previously defined in reference datasets from human middle temporal gyrus (MTG) and mouse primary visual cortex (VISp; single-nucleus or snRNA-seq in human, single-cell or scRNA-seq in mouse) (*21, 27*). With annotations from layer dissections as a guide, we identified 10 L1 t-types in human and 8 in mouse (Methods, Fig S1A). In UMAP projections of transcriptomic space (Fig. 1A), many human t-types formed separated clusters, with others clustered in groups of related t-types, while mouse L1 t-types showed more continuous variability. This contrast suggests stronger transcriptomic specialization in human L1, similar to supragranular excitatory neurons (*28*), and indicates that more robust groupings of L1 types into highly distinct transcriptomic subclasses can be delineated in human. We grouped related t-types into L1-focused transcriptomic subclasses by quantifying the pairwise distinctness of t-types in terms of a d’ separation of likelihoods (*23, 24*). This formed three subclasses, with three t-types remaining ungrouped (Fig. 1B). Expression of the inhibitory subclass markers *PAX6* and *LAMP5* (*27, 29*) and the t-type marker *MC4R* also closely matched these subclass boundaries (Fig. 1C).

**Fig. 1.**
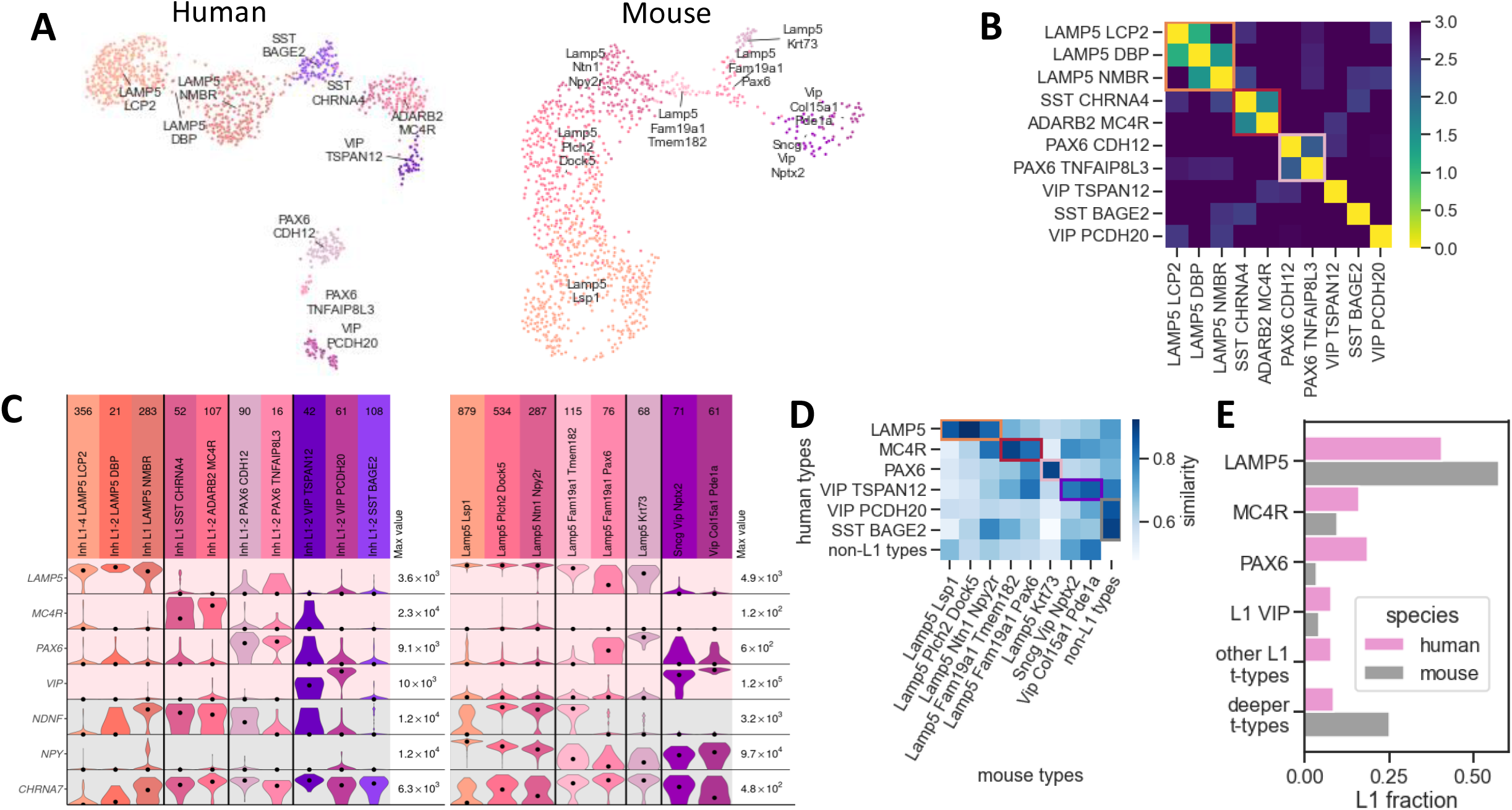
Single-nucleus RNA-seq demonstrates L1 diversity and provides a reference for patch-seq transcriptomic mapping. A. UMAP projections of human (left) and mouse (right) gene expression for L1 cell types (single neuron or nucleus RNA-seq)
B. Human t-types can be grouped into three subclasses based on transcriptomic similarity d’, with three ungrouped t-types remaining
C. L1 t-types are distinguished by expression of canonical and t-type-specific marker genes in human (left) and mouse (right). Pink background: human subclass markers, grey: classical mouse markers. Vertical lines divide group t-types by subclass. Violins show expression in log(CPM+1), normalized by gene for each species (maximal expression noted at right).
D. Mouse t-types are linked to homologous subclasses based on similarity in integrated transcriptomic space. Non-L1 t-types are excluded, with maximal similarity over all non-L1 types shown for reference.
E. Proportions of subclasses and unclassified t-types in L1, by species. Human proportions are from patch-seq cells with soma located in L1, mouse are from snRNA-seq reference restricted to L1 dissection, as mouse patch-seq was from targeted mouse lines which would bias proportions. Other L1 t-types refers to t-types in human L1 with no mouse homologue in L1. Deeper t-types refers to types found in L1 that do not meet the criteria for being a core L1 t-type. All comparisons significant at FDR-corrected p<0.05, one vs rest Fisher’s exact tests.

Human subclass marker genes did not clearly identify subclasses in mouse, which posed a challenge for cross-species comparison. Marker genes were either not expressed in any mouse L1 type (e.g. *MC4R*) or were expressed broadly and overlapped with other markers (e.g. *LAMP5*) (Fig. 1C top). Similarly, markers previously suggested for L1 subclasses in mouse (*19*) showed graded or complete lack of expression in human L1 (Fig 1C bottom).

Given the lack of conserved markers across species, we instead grouped mouse t-types for cross-species analysis by using cluster distances in an integrated transcriptomic space (*21*) (Fig S1C), identifying each mouse t-type with the most similar human subclass or ungrouped t-type (Fig 1D, Fig S1B). These matches formed four homology-driven subclasses (called subclasses hereafter) with proportions largely comparable across species (Fig 1E; PAX6 is the notable exception), named by the subclass marker genes in human. Two additional human L1 t-types (SST BAGE2 and VIP PCDH20) were excluded from cross-species L1 subclasses based on their homology to deeper t-types in mouse, suggesting a shift in some of the interneuron diversity across laminar boundaries between mouse and human (Fang et al., 2022). Reinforcing the validity of these subclass divisions in mouse, we noted likely matches to previous mouse L1 subclasses (*19*) based on marker gene expression (Fig 1C, Table S1): neurogliaform cells (*Npy+/Ndnf+*)to LAMP5, canopy cells (*Npy-/Ndnf+*) to MC4R, and α7 cells (*Ndnf-/Vip-/Chrna7+*) to PAX6 (*11*). However, uncertainty in these matches highlights the need for further confirmation based on morpho-electric properties.

To characterize morpho-electric and transcriptomic diversity across human L1 cell types, we used a previously established pipeline for high-throughput data acquisition and analysis (*26, 28*) to generate and release a comprehensive L1 patch-seq dataset. Human tissue was obtained from surgical samples and processed with standardized protocols; most samples originated from the MTG, along with smaller fractions in other temporal and frontal areas (Data S1). All cells were filtered for transcriptomic (n=420) and electrophysiological quality (n=252), and a subset of neurons (n=86) with sufficient cell labeling were imaged at high resolution and their dendritic and axonal morphologies were reconstructed.

We assigned transcriptomic cell types and subclasses to patch-seq samples using a “tree mapping” classifier, a decision tree based on the transcriptomic taxonomy structure (Methods) (*24*). Validating these assignments, we visualized t-type labels from patch-seq and reference datasets in a joint UMAP projection using alignment methods from the Seurat package (*30*) and found strong correspondence (Fig S1D). Additionally, marker genes used by the classifier showed strong correlation by t-type between patch-seq data and the snRNA-seq reference (Fig S1E).

Since patch-seq sampling was not uniform across cortical layers, we also measured the laminar distribution of L1 t-types using spatially resolved single-cell profiling of gene expression (multiplexed error-robust fluorescence in situ hybridization or MERFISH; Methods). These results were compatible with those from layer dissections of snRNA-seq, confirming that human L1 t-types are predominantly found in L1 or on the L1/L2 border, and demonstrating t-type-specific distributions across deeper layers and within L1 for certain types (Fig S1G, H).

### Morpho-electric diversity in human L1

Organizing the patch-seq dataset by transcriptomic subclass revealed the exceptionally diverse morphology and physiology of human L1 interneurons. Morphologically, subclasses were distinguished by vertical orientation of axons and dendrites, axon extent and shape, and dendrite branching (Figs 2A,D, S3; Data S2). Electrophysiologically, subclasses were distinguished by subthreshold properties such as sag (steady-state hyperpolarization reduced from transient peak) as well as several suprathreshold properties including firing rate, single action potential kinetics and adaptation of spike kinetics during trains of action potentials (Figs 2B, C; Data S2). Notably, spike adaptation properties showed a strong inverse relationship with sag across the dataset (Fig S2A). Sag is often mediated by HCN channels (*31*) and spike broadening by specific K+ channels (*32, 33*), so this finding may indicate a functional relationship between these channels in all subclasses of human L1 neurons.

**Fig. 2.**
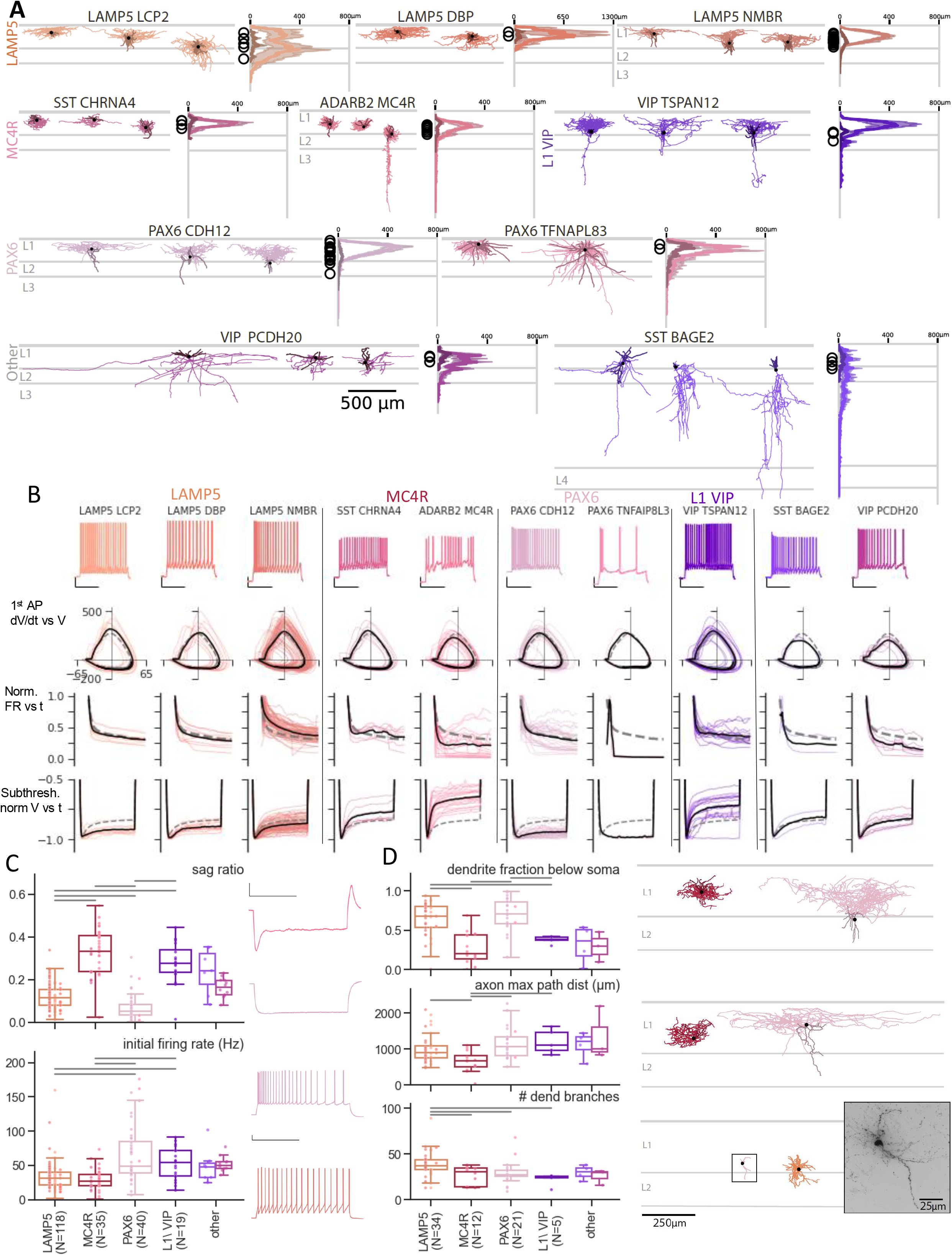
Human L1 transcriptomic subclasses are morpho-electrically diverse. (A): Example human morphologies for L1 t-types are displayed by subclass. Neurons are shown aligned to an average cortical template, with histograms to the right of the morphologies displaying average dendrite (darker color) and axon (lighter color) branch length by cortical depth for all reconstructed cells in L1 and L2 (shading shows +/-1 SD about mean, soma locations represented by black circles)
(B) Electrophysiology summary view by t-type and subclass. Top shows example spiking response at 40pA above rheobase (scalebar 20 mV, 0.5 s). Cell-by-cell summary traces shown below, with black t-type average, dashed dataset average, individual cells in color. Top to bottom: phase plane (dV/dt vs V) plot of first action potential; instantaneous firing rate normalized to peak; hyperpolarizing response normalized to peak.
(C-D) Electrophysiology and morphology features distinguishing L1 subclasses. Boxplots show subclass statistics (box marks quartiles, whiskers extend 1.5×IQR past box), with individual cells arranged horizontally by t-type. Significant pairwise comparisons marked above (FDR-corrected p<0.05, Dunn’s test post-hoc to KW test). Illustrative traces (electrophysiology, scalebar as in B) or layer-aligned reconstructions (morphology) shown for high and low values of each feature. Image inset shows that sparse dendrites in human PAX6 cells are not due to inability to resolve dendrites.

LAMP5 cells, the largest subclass, corresponded to the classical neurogliaform cell type (*34*), with highly branched, descending dendrites and horizontally elongated axons, either with a rectangular or triangular shape. Their electrophysiological phenotype was relatively undistinguished, with firing-rate adaptation and sag present but small. PAX6 cells had similar axons to LAMP5 cells, occasionally with descending branches, and sparser downward dendrites, along with minimal sag and high initial firing rate at the onset of response to depolarizing current injection. MC4R cells had extremely compact ball-like axonal arbors, along with the strongest sag; on this basis, they were tentatively identified as a match to the recently discovered ‘rosehip’ type (further characterized below) (*20*). L1 VIP (TSPAN12 t-type) cells had descending axon collaterals (*13*) with a consistent stellate-like dendrite morphology and high sag. The two cell types with no matching t-types within mouse L1, SST BAGE2 and VIP PCDH20, showed extremely diverse dendritic and axonal structure, often with significant horizontal or descending axon branches – even avoiding L1 entirely in the case of some BAGE2 cells. These t-types were more uniform electrophysiologically, with relatively small spikes, high adaptation and sag, but were sparsely sampled and thus difficult to fully characterize – in particular, SST BAGE2 cells were sampled at a significantly lower rate than in our snRNA-seq dataset (Fig S1F).

In a few instances, we also observed differentiation between t-types within the same subclass. Within the LAMP5 subclass, sag and adaptation decreased from the NMBR t-type to DBP to LCP2 t-types. While other LAMP5 t-types were mostly restricted to L1 and superficial L2, the LCP2 t-type was found distributed across all cortical layers, with axonal arbors becoming less elongated and overlapping less with dendritic arbors for deeper cells (Fig. S1G,H; S2B). PAX6 cells were distinguished by whether the initial high-frequency firing formed a discrete burst (TNFAIP8L3) or continuously adapted (CDH12). The two MC4R t-types were distinguished by the magnitude of sag and irregularity of firing.

Given the potential for the observed neuronal diversity to be determined in part by diversity in tissue donor characteristics, we tested all morpho-electric features for effects of donor medical condition, sex, and age (Fig S5A,B; Data S3). Most effects were small and in features not linked to L1 diversity, with the notable exception of higher dendritic branching in cells from tumor patients compared to epilepsy patients (Fig SB5; this result was not explained by brain area or subclass).

### Cross-species differences in L1

Evolutionary expansion of L2/3 in primates was previously linked to changes in cytoarchitecture, including thinning out of cell density and increased soma size, accompanying the specialization of pyramidal cell types (*28*). Considering this, we first quantified cytoarchitecture differences in L1 of human tissue samples (NeuN-stained slices from patch-seq tissue blocks) compared to mouse samples (Fig. 3A). In contrast to cross species differences in L2/3, human L1 was thicker, but with smaller, denser cell bodies that were more evenly distributed across the layer compared with mouse, agreeing with prior observations (*35*).

**Fig. 3.**
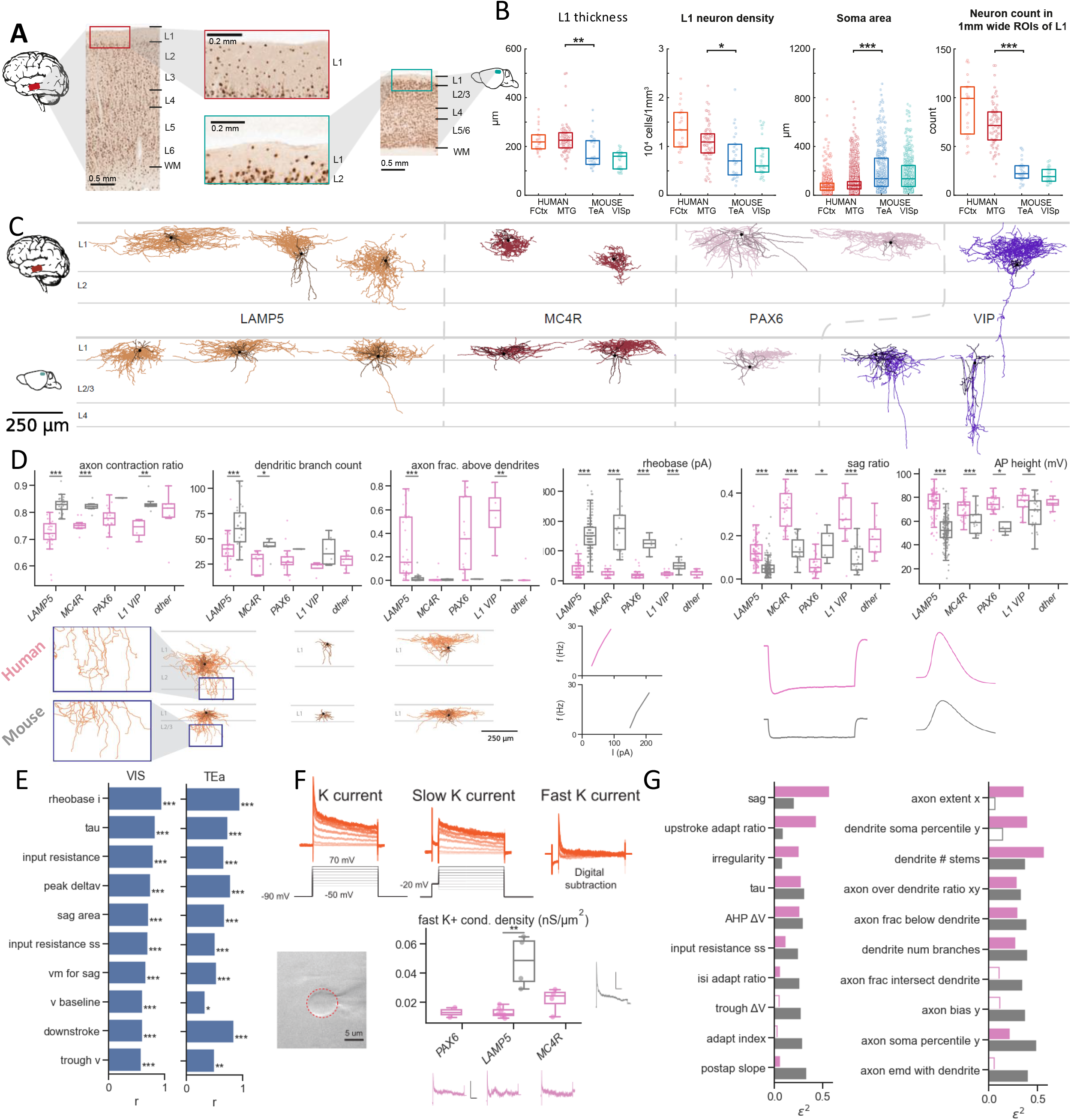
Comparison of human and mouse L1. A. Examples of NeuN labelling of neurons in human MTG and mouse VISp.
B. Comparisons of mouse versus human L1 thickness, neuron density, soma area and neuron count in 1mm wide ROIs of L1. Metrics plotted per ROI for L1 thickness, density and neuron count, and per cell for soma area. Boxplots show quartiles, stars indicate post-hoc Dunn’s test results at p<[0.05, 0.01, 0.001] (calculated for MTG vs TEa only).
C. Example layer-aligned morphologies from mouse and human L1 subclasses. One example shown from each t-type, scalebar for both species.
D. Morphology (left) and electrophysiology (right) features with differences between human and mouse L1 cells. For features with a species-subclass interaction (2-way ANOVA on ranks, p<0.05 FDR-corrected), stars indicate post-hoc Dunn’s test results at p<[0.05, 0.01, 0.001]. Representative examples shown below each plot (L to R: layer-aligned reconstructions, AP frequency as a function of current injection, response to hyperpolarizing current, first action potential).
E. Electrophysiology feature differences between human L1 and mouse VISp L1 (left) largely hold when tested against mouse TEa (right). Features selected by largest effect size against TEa (MW r, rank-biserial corr). Stars indicate significance (FDR-corrected MW test, p<[0.05, 0.01, 0.001]).
F. Nucleated patch recordings revealed higher A-type K+ conductance in mouse. Example traces show voltage commands (black) and recorded currents (orange) from measurement protocol (top),along with example soma size measurement. Boxplots show fast conductance density in both species, with example traces shown for each group (scalebars 400pA/200ms)
G. Different sets of features distinguish human and mouse subclasses. Bars show size of subclass effect (eta^2 from 1-way ANOVA), with features ranked by the difference between human and mouse effects. Unfilled bars indicate p>0.05 (FDR-corrected F test).

To study morpho-electric differences in L1, we compiled a comparison patch-seq dataset from mouse L1 neurons (n=531, 76 with morphology) consisting of previously published data from a cross-layer analysis of interneurons in VISp (*24*) and additional recordings in L1 and L2/3 of visual cortex and the temporal association area (TEa; held out as a validation set as a region that is often compared to human MTG (*36, 37*). Despite the differences in L1 cytoarchitecture, morphologies of L1 neurons generally showed remarkable similarity across species when comparing across matched homology-driven subclasses (Fig 3C, S3, S4; Data S4). Although human neurons were slightly larger in horizontal extent, no differences were observed in vertical dimensions or dendritic diameter (Fig. S2C). No mouse L1 neurons had morphologies resembling the unmatched human L1 t-types (VIP PCDH20 and SST BAGE12, homologous to deeper mouse t-types), suggesting that these cell types are unique to human L1. Mouse VIP cells had descending axon branches as in human VIP cells, but with greater variability of structure. Mouse LAMP5 cells had dense neurogliaform-like axonal arbors, confirming their match via *Ndnf/Npy* expression. Unlike human axons, mouse axons rarely extended above dendrites (Fig. 3D left), perhaps reflecting sublaminar structure found only in the thicker human L1. Human neurites were also structured differently, with smaller contraction ratios (higher tortuosity) compared to the straighter but more heavily branched mouse dendrites – possibly an adaptation to the higher cell density.

As in previous studies, electrophysiological properties showed stronger differences across species (*38*). Mouse cells had no sag, much broader spikes, and higher rheobase across at least 3 of 4 matched subclasses (Fig 3D; Data S4). We replicated these findings in a comparison between MTG and a smaller TEa dataset, verifying that cross species differences were not due to regional differences between MTG and VISp (Fig 3E). Proportions of L1 t-types also varied little across brain regions in mouse and human single neuron/cell RNAseq reference datasets (Fig S1G). We explored dependence of L1 interneuron morpho-electric properties on brain region within our human data and found moderate effects on a set of features including dendrite extent and input resistance (Fig S5D). MTG L1 cells had larger dendrite extent and lower input resistance (closer to mouse cells), suggesting again that the cross-species differences were not inflated by different brain regions sampled.

To investigate causal factors underlying cross-species electrophysiology differences, we looked for correlated differences in ion channel gene expression and morphology features compared against membrane properties in the patch-seq dataset. The increased branching of mouse cells could affect input resistance if branching is proximal to the soma, by increasing the effective membrane area for leak conductance. We found a higher peak of total dendrite cross-sectional area at ^~^50 um from the soma as well as slightly higher total volume in mouse cells (Fig S2F) supporting this explanation. Differences in spike shape and threshold could be explained by potassium channel differences, along with related features like rheobase and delayed spiking. Indeed, the expression of genes (*KCND2, KCND3*, and *KCNH7*) associated with fast inactivating, A-type K+ channels (Kv4.2, Kv4.3 and the ERG3 channel Kv11.3 (*39, 40*)) was higher in mouse neurons and was correlated with several action potential features (Fig S2D). To test for corresponding differences in K+ channel conductance, we measured macroscopic currents in nucleated patches following whole-cell recording in a subset of cells. Compared with human neurons, mouse neurons showed much higher A-type K+ conductance but comparable slow inactivating (D-type) conductance (Fig 3F). Considering blocking Kv4 channels in mouse neurogliaform cells decreases AP threshold and latency of first AP onset (*41*), these differences in A-type K+ channel conductance, along with lack of Kv1.1 expression, may contribute to the lack of late spiking observed in human L1 neurogliaform cells as well (*42*).

Finally, we asked whether the strong morpho-electric variability observed between human L1 subclasses is also present in L1 of mouse neocortex. Ranking electrophysiology and morphology features by the amount of variability between subclasses they explain, we found that the two species had a similar amount of variability (number of significantly different features and their effect size) but varied along different sets of features (Fig 3G). The most distinct features in human, like sag and spike shape adaptation, showed little variability in mouse, and unlike in human, mouse subclasses varied physiologically in ISI adaptation and spike after-hyperpolarization (AHP) properties, and morphologically in relative vertical positioning of the axonal arbor (most features largely driven by L1 VIP subclass: Fig. S2; S4).

### Distinctive neuronal phenotypes in human L1

Despite the quantitative similarity in L1 heterogeneity across species, we noted two particularly distinctive phenotypes found in human L1 only. The MC4R rosehip cells and the bursting PAX6 TNFAIP8L3 t-type were both qualitatively distinct from other L1 types, whereas morpho-electric variability in mouse was more continuous, as we also observed with transcriptomic variability (Fig. 1A). To further highlight this contrast, we investigated each of these highly distinctive types in turn by quantifying the distinctive morpho-electric features and marker genes, then searching for comparable cells in the mouse L1 dataset.

### Rosehip cells

The MC4R subclass, putative rosehip cells, comprises two transcriptomically similar t-types, SST CHRNA4 and ADARB2 MC4R, both highly distinct from other L1 types including the LAMP5 LCP2 t-type originally identified with the rosehip phenotype. MC4R morphologies were all confirmed to qualitatively match the distinctive rosehip axonal structure and boutons (Fig. 4A, S3), and were quantitatively distinct from other L1 types in terms of maximum axonal path distance and branch frequency (Fig. 4B). We also noted two examples of MC4R cells (both within the ADARB2 MC4R t-type) with elaborate descending axons reaching the lower half of L3, and confirmed that the characteristic large, dense axonal boutons were visible on both the central axonal arbor and descending axons when present (Fig 4A right). Electrophysiologically, both t-types that comprise the MC4R subclass showed strong sag, but only the ADARB2 MC4R t-type showed irregular firing (*42*). Cells in the ADARB2 MC4R t-type also had somas and axons localized near the L1/L2 border (Fig S1H, S6A). We explored the expression of genes related to neuron physiology (ion channels and GPCRs) and found markers that both distinguish the entire rosehip subclass from the rest of L1, including *HTR1F* (5HT receptor 1F), along with markers distinguishing the rosehip subtypes: *GRM5* (metabotropic glutamate receptor 5) and *RELN* (Reelin) showed lower expression in ADARB2 MC4R neurons in particular (Fig 4C). Together, these differences in gene expression and physiology indicate that there are distinct rosehip neuron subtypes within human L1.

**Fig. 4.**
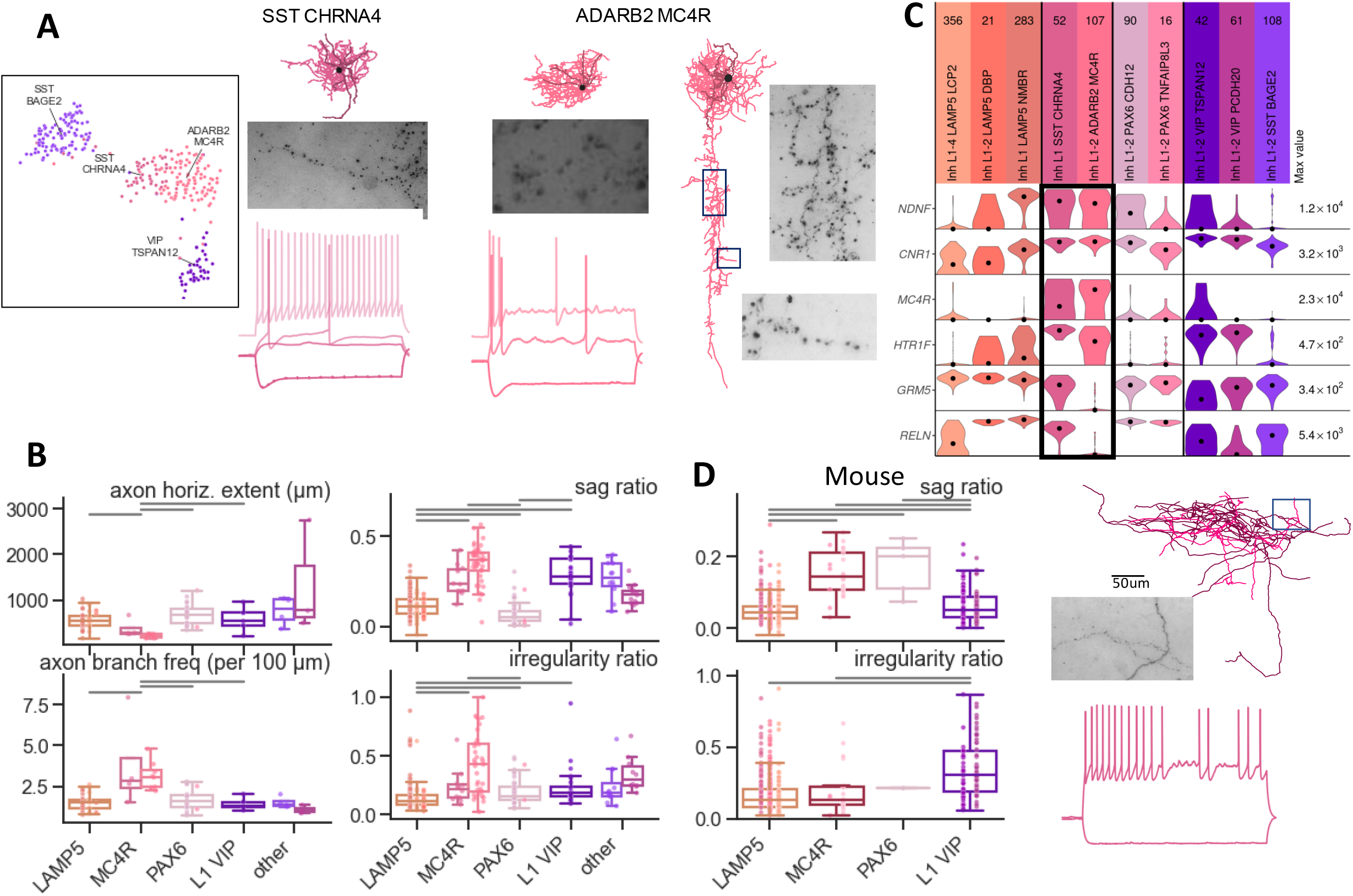
MC4R rosehip cells. A. Example rosehip cells from MC4R subclass: UMAP projection of transcriptomic data from MC4R and nearby subclasses (left); morphologies with insets showing 63x MIP images of axonal boutons (top right; notice lack of rosehip boutons in the mouse neuron in panel D); example electrophysiology traces (bottom; hyperpolarization near −100 mV, depolarization below rheobase, spiking at rheobase and 40 pA above)
B. Electrophysiology and morphology features distinguishing MC4R t-types (highlighted). Boxplots show t-type statistics (box marks quartiles, whiskers extend 1.5×IQR past box),
C. Gene expression of MC4R subclass (highlighted) and other L1 t-types, for between- and within-subclass marker genes (snRNA-seq). Violins show expression in log(CPM+1), normalized by gene (maximal expression noted at right).
D. Mouse L1 included cells with moderate sag and irregular firing, but no cells with both properties (boxplots as in C). Example morphology and electrophysiology shown for mouse L1 LAMP5 cell with highly irregular firing, but lack of rosehip-like morphology.

In mouse L1, there were no cell types observed with the morphological signatures consistent with human rosehip cells (Fig S4) and only partial matches to the established electrophysiological signatures: the homologous MC4R subclass had moderate sag but no irregular spiking, while irregular spiking resembling the ADARB2 MC4R rosehip t-type was present only in a subset of LAMP5 cells without other rosehip-like features (Fig 4D). The mouse MC4R subclass was, however, the closest match to the marker gene signature defining canopy cells (*Npy-/Ndnf*+). Some morpho-electric characteristics of canopy cells, including wide dendritic extent and moderate sag, were matched by mouse MC4R cells, but notably not the characteristic L1a-dominant axon. This defining feature was observed only in a subset of the Lamp5 Plch2 Dock5 t-type (Fig S6A-B) which do not match canopy marker genes (*Npy*+). The mouse MC4R subclass thus does not have clear rosehip phenotypes nor is it a clear match to the canopy type, but rather seems to be a subset of canopy-like cells with distinct transcriptomic properties partially aligned with the human MC4R subclass.

### Bursting PAX6 TNFAIP8L3 cells

The other highly distinctive firing pattern we noted in human L1 was in the PAX6 TNFAIP8L3 t-type, which fired in high-frequency bursts at the onset of stimulation, followed by quiescence or regular firing at higher stimulus amplitudes. Spiking and dendritic structure were highly distinct between this t-type and the neighboring PAX6 CDH12 t-type, despite some similarity of axonal structure and subthreshold electrophysiology (Fig. 5A). Both the initial firing rate at rheobase and the after-depolarization (ADP) following the final spike quantitatively distinguished PAX6 TNFAIP8L3 cells from all other L1 cells (Fig. 5B), as did the number of dendritic branches and large horizontal dendritic extent, over 550 microns wide.

**Fig. 5.**
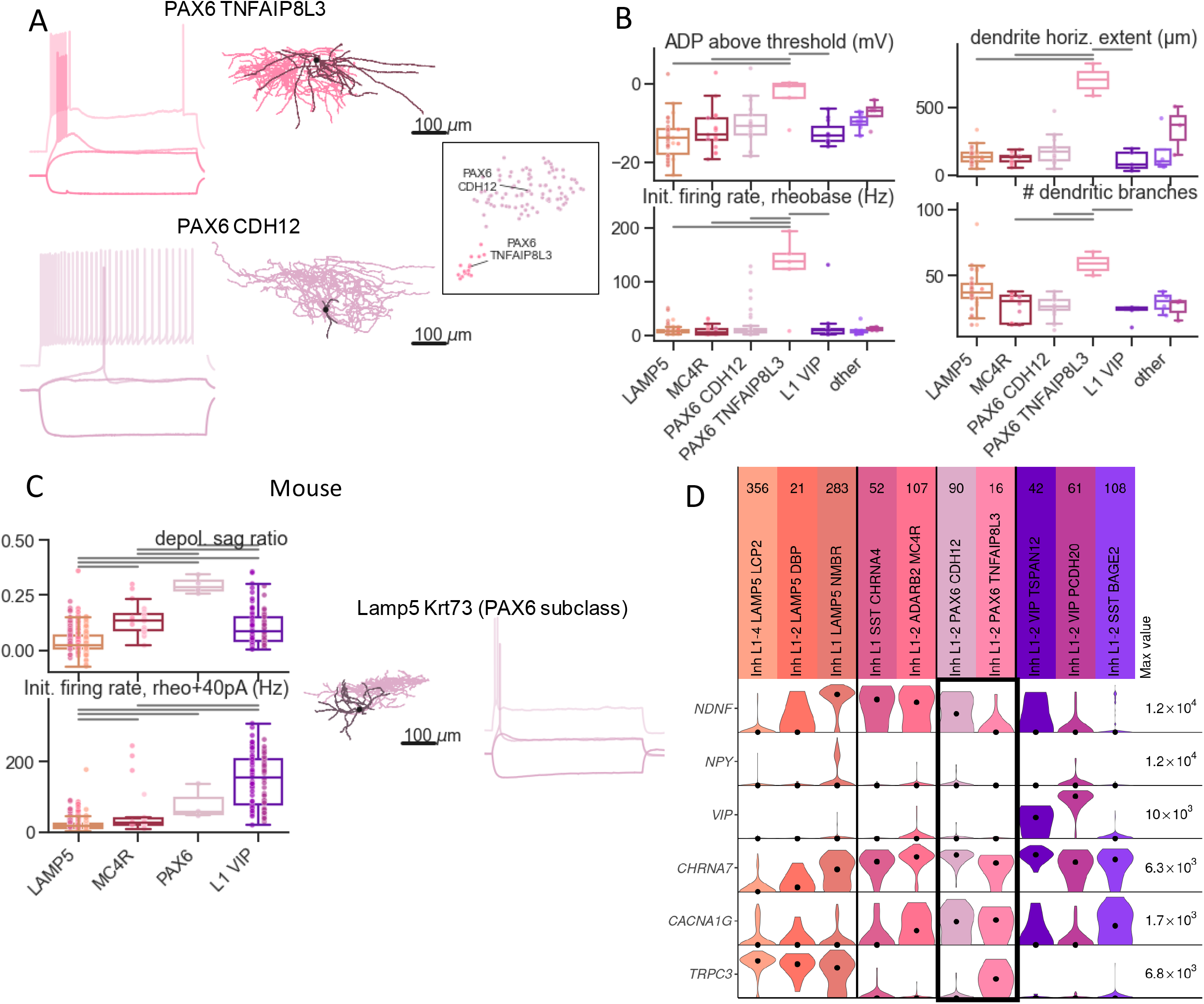
Burst spiking PAX6 TNFAIP8L3 cells. A. Reconstructed morphologies and example electrophysiology for bursting and non-bursting PAX6 t-types (TNFAIP8L3 and CDH12) (hyperpolarization near −100 mV, depolarization below rheobase, spiking at rheobase and 40 pA above). Inset shows UMAP projection of transcriptomic data from PAX6 subclass.
B. Electrophysiology and morphology features distinguishing PAX6 TNFAIP8L3t-type (PAX6 subclass highlighted). Boxplots show t-type statistics (box marks quartiles, whiskers extend 1.5×IQR past box),
C. Mouse L1 includes cells with initial doublet firing, but without longer bursts or long dendrites. Example morphology and electrophysiology shown from PAX6 subclass (Lamp5 Krt73 t-type) and *Pax6+* cell in Lamp5 Fam19a1 Pax6 t-type. Depolarizing sag ratio is the normalized size of the hump at stimulus onset just below rheobase.
D. Gene expression of human PAX6 subclass (highlighted) and other L1 t-types, for alpha7 type and bursting-related marker genes (snRNA-seq). Violins show expression in log(CPM+1), normalized by gene (maximal expression noted at right).

In mouse L1, the homologous PAX6 subclass was extremely rare, comprising only a few cells in the Lamp5 Krt73 t-type. These cells tended to fire in doublets at stimulus onset rather than a full burst, sometimes followed by a delayed ADP (Fig 5C). Some cells in the Lamp5 Fam19a1 Pax6 t-type also showed this firing pattern (Fig S6D), likely the same subset that align transcriptomically to the human PAX6 subclass (Fig 1C). Mouse doublet-firing cells also showed a depolarizing ‘hump’ for current injection just below rheobase, which together with the marker gene signature (*Ndnf-/Vip-/Chrna7+*) matches these cells with the mouse α7 type, a type previously defined in mouse by these physiological/gene features (*19*). This hump was suggested to indicate activation of T-type calcium channels, likely the same mechanism underlying the bursting in human cells (*42*). Bursting was also previously noted in a subset of mouse “Single Bouquet Cells” (SBC) (*25*), a group defined by loose morphological criteria. This class likely overlaps with the doublet-firing t-types (*43*), suggesting they may burst under different conditions.

Using these insights from the cross-species alignment, we explored the expression of related genes in the human PAX6 t-types (Fig. 5C). Both t-types matched the α7 marker gene signature (along with the SST BAGE2 t-type; *Ndnf-/Vip-/Chrna7+*), and strongly expressed the T-type calcium channel alpha subunit *CACNA1G*, highlighting T-type calcium channels as a potential factor in the burst and doublet firing across species. Given the lack of bursting in PAX6 CDH12 cells, other ion channel genes differentially expressed between the two human PAX6 t-types likely also play a role, including *TRPC3*, a non-specific cation channel that can regulate resting membrane potential (*44*).

### Cross-modality relationships of L1 subclasses and t-types

Given the multiple observations of distinctness between human types contrasted with continuous variation between mouse types, we explored this contrast more comprehensively by defining a common quantitative framework for distinctness across modalities. We generalized the d’ metric used for transcriptomic distinctness (*23, 24*), quantifying the performance of classifiers trained to distinguish pairs of t-types based on electrophysiology and morphology features. The resulting t-type similarity matrices (Fig 6A) showed comparable subclass structure in both electrophysiology and transcriptomics, with smaller d’ values within subclass blocks and higher values outside. Notably, d’ metrics were highly correlated between modalities, demonstrating that cell types with distinctive transcriptomes have similarly distinctive electrophysiological properties (Pearson r=0.64, p=0.00021; Fig 6B). The single within-subclass pair with a high d’ was LAMP5 NMBR and LAMP5 LCP2, which sit at opposite ends of the LAMP5 continuum. We also calculated d’ similarity matrices at the subclass level to allow comparison between species in all three modalities (Fig 6C). These results confirmed the generally higher distinctness of subclasses in human, while in the mouse only the VIP subclass was highly distinct in all modalities.

**Fig. 6.**
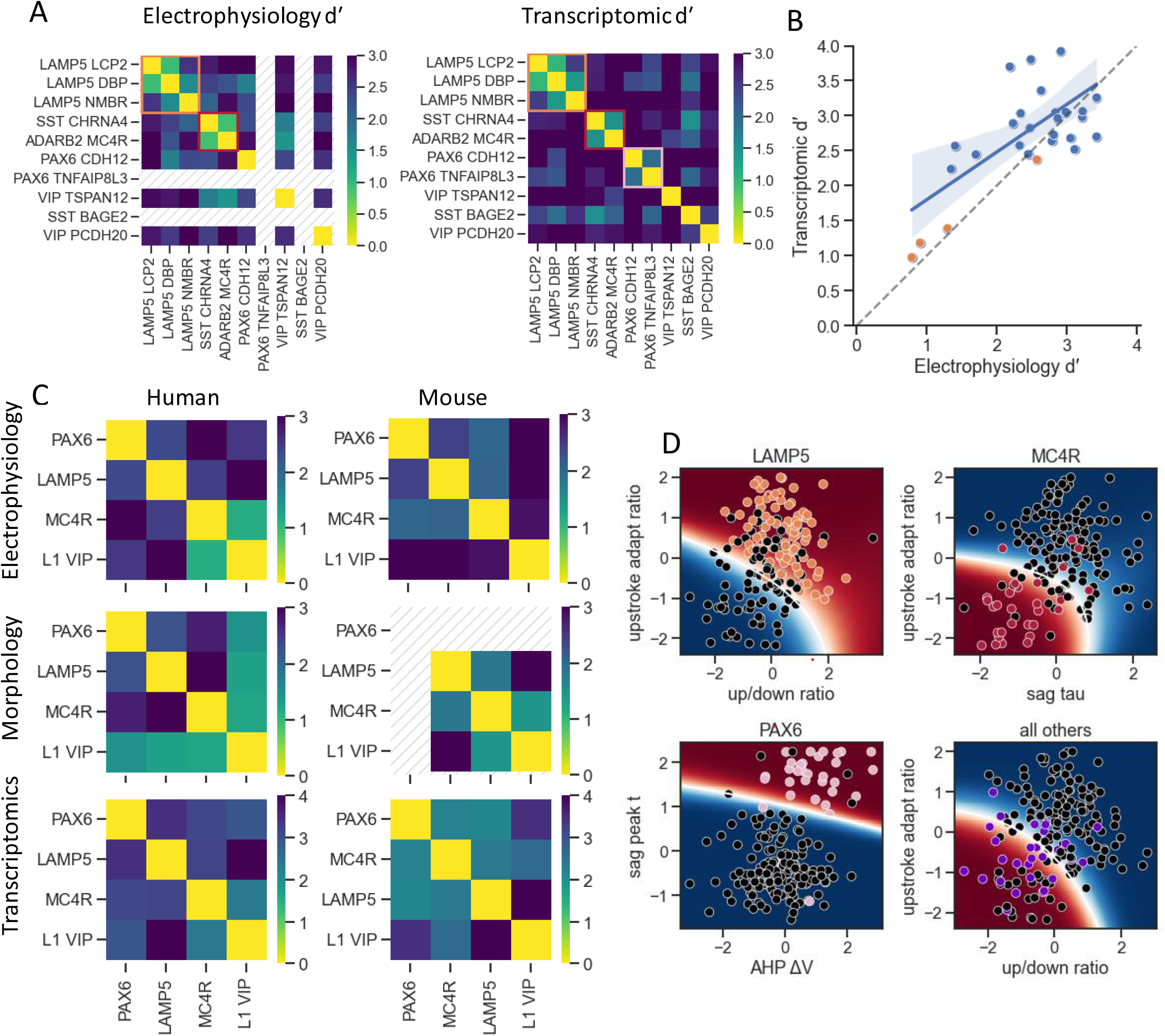
Quantifying distinctness of L1 t-types and cross-modality structure. A. Pairwise similarity matrices of L1 t-types, from classifiers using electrophysiology (left) and gene expression (right). D’ (d-prime) is a metric of separation of distribution means, scaled relative to the standard deviation. Groups with N<10 excluded (hatched area).
B. Correlation of pairwise d’ values between transcriptomic and electrophysiology feature spaces shows similarity of dataset structure across modalities. Shading shows bootstrapped 95% CI of regression. Smaller d’ for within-subclass pairs (orange) shows subclass structure.
C. Similarity matrices of L1 subclasses across species and data modality. Groups with N<10 excluded.
D. L1 subclasses cluster separately in electrophysiology subspaces. Points show all L1 neurons, with the subclass of interest in color. Background color shows cluster membership likelihoods from 2-cluster Gaussian mixture model trained on unlabeled data. F1 scores: LAMP5 0.81, MC4R 0.69, PAX6 0.89, all others 0.5 (L1 VIP and ungrouped t-types). All features normalized, Yeo-Johnson transform applied to approach Gaussian distribution.

To visualize the subclass-level distinctness in terms of specific electrophysiology features, we found the pair of features that most distinguish each subclass and showed that clusters defined by these features closely match the transcriptomic subclass boundaries (Fig 6D). We also tested the effectiveness of assigning subclass labels to neurons based on the full electrophysiology feature set. A multi-class classifier evaluated by cross-validation on the primary dataset had 82% accuracy balanced across subclasses (Fig S7A). To mimic the out-of-sample issues that could be encountered for future L1 datasets collected under different conditions, we also tested classifier performance on data held out of our primary analysis due to equipment and protocol differences. After excluding features for which the distributions strongly differed from the primary dataset, we found comparable classification performance (81%, Fig. S7B), reinforcing the utility of the human L1 subclasses for understanding L1 variability even in the absence of transcriptomic information to assign subclass identity.

## Discussion

### Summary

Using patch-seq, we identified a coherent view of human L1 interneurons in which neuronal subclasses defined by transcriptomic distinctness have similarly distinct morpho-electric phenotypes. Two human cell types emerged with especially distinct phenotypes that were not matched in their putative homologues in mouse: the compact, high-sag MC4R ‘rosehip’ subclass and the large, burst-spiking PAX6 TNFAIPL83 t-type. Although mouse L1 neurons had a similar range of diversity in most features, human L1 neurons spanned a wider range of sizes and generally showed stronger distinctions between subclasses across modalities, similar to the specialization observed in human L2-3 pyramidal cells (*28*). Human and mouse neurons also showed consistent differences in certain morphological and physiological properties across all subclasses, despite a general similarity in cell size. These results indicate a general conservation of L1 inhibitory neuron diversity, but with significant specializations in cell properties and subclass/cell-type proportions, likely leading to differences in the regulation of higher order input to the human cortical circuit.

### Categorizing L1 neuron types

Our results in mouse support previous classification schemes consisting of four primary types in L1 but also clarify a need for precise data-driven criteria for those types. The near-alignment of our homology-driven subclasses and cell types based on single marker genes (*19*) provides additional support to the validity of those types, but identification of marker-gene-defined types in our data was in some instances ambiguous or lacked alignment with criteria in other modalities. Similarly, other coarse single-modality cell-type distinctions in L1 (late spiking vs non late spiking, NGFC vs SBC) likely grouped multiple distinct subclasses and shifted the exact boundaries. In part, the consensus view of cell type diversity in L1 should consider a larger role for continuous variability. A continuous transition between *Ndnf+/Npy+* neurogliaform cells and *Ndnf+/Npy*-canopy cells was previously noted (*24*); we additionally observed continuity between PAX6 (putative α7 L1 neuron subset) and MC4R (putative L1 canopy subset) types, thus helping to explain observations of distinct phenotypes spanning subclass or t-type boundaries in mouse L1 (canopy-like L1a axons in some LAMP5 cells, α7-like doublet spiking in some MC4R cells). The ambiguity of certain subclasses in mouse also highlights the remarkable alignment of transcriptomic subclasses in human L1 with electrophysiological and morphological divisions, likely due to the increased distinctiveness of human subclasses. A mouse transcriptomic classification better informed by cross-species or cross-modality insights might achieve similar alignment and more precisely unify different views of mouse L1 types.

Our work demonstrates the benefits of detailed transcriptomic data over small sets of genetic markers for both accurately characterizing cell type divisions and establishing cross-species homologies that facilitate comparative analysis. Within species, reliance on marker genes can overstate the distinctness of cell types, as with the NGFC/canopy distinction, or even lead to misidentification, as with the original description of human L1 rosehip cells (*20*). Across species, the lack of conserved L1 markers was striking and perhaps unique to layer 1. Even using the full transcriptome, previous work on this homology found variable results for L1 types with different brain areas and methods (*21*) – we chose to quantify similarity of t-types across species in a way that better captured ambiguity and found strong matches for some types and weaker for others. Ambiguous matches may indicate areas of evolutionary change which could be illuminated by comparative or developmental analyses of additional species phylogenetically related to mouse and human. For instance, the weak cross-species transcriptomic similarity of the MC4R subclass (Fig 1C) and lack of phenotypic similarity together suggest that human and mouse MC4R cells could represent distinct innovations in each taxon, rather than a true homology.

In human L1, the strong alignment of subclass distinctions across modalities suggests that cells can be classified using only morphology or electrophysiology with reasonable accuracy. Condition-dependent variation of certain electrophysiology features can present a challenge to this approach though, especially for classifications relying on small numbers of features with especially strong qualitative variability. We failed to observe two electrophysiological phenotypes that had been noted in past work: late spiking in human NGFCs (*38*), and full bursting in mouse SBCs (*25*), qualitative features for which conflicting observations have also been reported (*19, 45*). While potential contributing factors are numerous (age differences, donor characteristics, recording conditions including internal solutions, temperature and equipment differences), we showed that for classification with a large feature set, identifying and excluding affected features can rescue reliable performance.

### Cell types, evolution, and function

Given the dramatic evolutionary expansion of layers 2 and 3 in primate neocortex and the known role of L1 inhibition in regulating dendritic integration in L2/3 pyramidal cells dendrites, it is likely that some of the increased complexity of human L1 types evolved to support the increasingly complex role of L2/3 in the cortical circuit (*28, 46, 47*) – new types of pyramidal cells might necessitate new types of dendritic inhibition. Similarly, the L1 circuit might also require adaptation to the decreased proportions in primates of L5 extratelencephalic pyramidal neurons, the prominent apical dendrites of which are targeted by L1 inhibition in rodents (*21, 22, 48*). Rosehip cells were previously shown to inhibit pyramidal cell apical dendrite shafts in L2/3 (*20*); the rosehip subtypes, with distinct electrophysiology could plausibly perform similar but distinct inhibitory functions or selectively modulate different pyramidal neuron subtypes in L2/3 (*28*). The presence of distinct VIP types in human L1 which are located in deeper layers in mouse could also have circuit implications, considering the selectivity and sublaminar segregation of thalamic projections to *Vip+* vs *Ndnf+* cells in mouse L1 (*49*)

While neurogliaform cell properties were largely conserved, cross-species differences may indicate varying demands on the types of GABAergic (e.g. synaptic and extra-synaptic) and gap-junction mediated transmission they provide, which depend critically on the spatial properties of the axonal arbor (*34, 50, 51*). LAMP5 axonal arbors were larger in human than mouse (^~^1.2×), but this increase was much less than the 1.6× increase in pyramidal cell apical dendrite extent in L1. The accompanying increase in soma density in human L1 could enable the conservation of the ‘blanket’ inhibitory function while also permitting some increased spatial/topographic selectivity. Neurogliaform circuit connectivity has been shown to be both tightly controlled (*52, 53*) and to exert strong effects on pyramidal cell sensory processing (*1, 6*). Neurogliaform cell density changes are thus likely to have either a direct or a compensatory function, perhaps linked to broader changes in excitatory to inhibitory cell ratios between mouse and human (*21, 22, 54*).

The strong bursting dynamics and distinctive morphology of the PAX6 TNFAIP8L3 t-type clearly point to a unique functional role. Their extended dendrites are well positioned to integrate local pyramidal cell inputs across a broad spatial footprint and long-range axonal inputs across topographic boundaries, and the bursting would provide a strong immediate activation in response to strong or coincidental input. The clear identification of a cross-species homology for the PAX6 subclass can aid in deciphering its function, combining functional insights from manipulating mouse cells with indirect insights from the more distinctive morphologies of human cells. Conflicting connectivity patterns have been observed for coarser cell types that likely include some PAX6 cells: mouse α7 cells synapse onto nearby L2 pyramidal neurons (*8*), while rat SBCs synapse onto L2/3 interneurons (*55*). More focused investigation of PAX6 cell connectivity is thus needed to illuminate the function of this subclass, and in turn the functional implications of specialization within this subclass in human L1.

Much experimental evidence has documented the importance of neuromodulatory control on L1 function (*38, 56*–*58*). Linking distinct subclasses in L1 to detailed gene expression data can help to suggest refined hypotheses for cell-type specific neuromodulation. In particular, human MC4R cells uniquely expressed several modulatory receptor genes (Fig S8A), including the melanocortin receptor *MC4R*, which plays a role in energy homeostasis in hypothalamus (*59*), serotonin receptor *HTR1F* and metabotropic glutamate receptor *GRM1* (along with differential expression of *GRM5* between MC4R t-types). Cholinergic activation of L1 cells (*56*), suggested to control attention, may also differentially modulate MC4R cells relative to other L1 types based on their stronger *CHRNA6* and *CHRNA4* expression.

These observations highlight the wealth of hypotheses that can be generated from the comprehensive human L1 patch-seq dataset reported here. Our analysis provides new tools for the classification of L1 diversity in both human and mouse, new insights into functional relationships underlying physiological differences, and the clear identification of subclasses and subtypes that are likely to be of particular interest in functional studies, all important steps for deciphering the function of this enigmatic layer of neocortex. This approach also represents a roadmap for annotating functionally related properties onto transcriptomically-defined cell type taxonomies that could be applied across the primate brain – a crucial step towards linking cell type diversity to functional diversity within a neural circuit.

## Author contributions

Writing – original draft: TC, RD, BK

Writing – review & editing: TC, RD, BK, NAG, JAM, GM, AM, AG, JT, SS, EL

Conceptualization: TC, RD, BK, KC, JT, EL

Methodology: TC, RD, BK, JC, NAG, BRL, RM, JAM, GM, JB, MK, NWG, BL, TJ

Resources: TC, ND, TE, AG, MM, JN, PB, CC, RGE, MF, EJM, BG, RPG, JSH, CDK, ALK, JGO, AP, JR, DLS

Investigation: TC, RD, BK, JC, NAG, BRL, RM, JAM, GM, AM, LA, KB, TEB, JB, DB, KB, EAC, RE, AG, EG, JG, KH, TSH, DH, NJ, LK, AKK, MK, GL, BL, MM, MM, DM, NM, LN, VO, ZP, AP, LP, RR, MR, JS, IS, MT, MT, JT, SV, GW, JW, ZY PB, CC, RGE, MF, BG, RPG, JSH, TJ, CDK, ALK, JGO, AP, JR, DLS, KS, SAS, BT, JT, JW, CPJdK, HM, GT, EL

Visualization: TC, RD, BK, JC, NAG, BRL, RM, AM

Funding: NAG, GM, CPJdK, HM, GT, EL

Project administration: LK, JN, SMS, DV, LE

Supervision: RD, BK, BRL, MM, CK, KS, SAS, BT, JT, JW, CPJdK, HM, GT, HZ, EL

Software: TC, TB, TJ

## Data and materials availability

The primary datasets used in this study are hosted at the archives DANDI (dandiarchive.org, electrophysiology), BIL (brainimagelibrary.org, imaging and reconstructions) and NeMO (nemoarchive.org,transcriptomics). These datasets are cataloged for easy access at brain-map.org: knowledge.brain-map.org/data/97XR43ZTYJ2CQED8YA6/collections for human data, and knowledge.brain-map.org/data/1HEYEW7GMUKWIQW37BO/collections for previously published mouse visual cortex data.

Custom analysis and visualization code, along with intermediate datasets (extracted features) are hosted at github.com/AllenInstitute/patchseq_human_L1. Other software libraries essential to the analysis are publicly available at github.com/AllenInstitute/ipfx (electrophysiology), github.com/AllenInstitute/neuron_morphology and github.com/AllenInstitute/skeleton_keys (morphology), and github.com/AllenInstitute/scrattch/ (transcriptomics).

## Supplementary Materials

Materials and Methods

Figs. S1 to S8

Table S1

References (*60–74*)

Data S1 to S4 [Donor characteristics; Human subclass difference statistics; Metadata effect statistics; Cross-species difference statistics]

### Materials and Methods

Detailed descriptions of patch-seq data collection methods in the form of technical white papers can also be found under ‘Documentation’ at http://celltypes.brain-map.org.

#### Human tissue acquisition

Surgical specimens were obtained from local hospitals (Seattle - Harborview Medical Center, Swedish Medical Center and University of Washington Medical Center; Amsterdam - Vrije Universiteit Medical Center; Szeged – Department of Neurosurgery, University of Szeged) in collaboration with local neurosurgeons. Data included in this study were obtained from neurosurgical tissue resections for the treatment of refractory temporal lobe epilepsy, hydrocephalus or deep brain tumor (Data S1). All patients provided informed consent and experimental procedures were approved by hospital institute review boards before commencing the study. Tissue was placed in slicing artificial cerebral spinal fluid (ACSF) as soon as possible following resection. Slicing ACSF comprised (in mM): 92 *N*-methyl-D-glucamine chloride (NMDG-Cl), 2.5 KCl, 1.2 NaH_2_PO_4_, 30 NaHCO_3_, 20 4-(2-hydroxyethyl)-1-piperazineethanesulfonic acid (HEPES), 25 D-glucose, 2 thiourea, 5 sodium-L-ascorbate, 3 sodium pyruvate, 0.5 CaCl_2_.4H_2_O and 10 MgSO_4_.7H_2_O. Before use, the solution was equilibrated with 95% O_2_, 5% CO_2_ and the pH was adjusted to 7.3-7.4 by addition of 5N HCl solution. Osmolality was verified to be between 295–310 mOsm kg^-1^. Human surgical tissue specimens were immediately transported (10–35 min) from the hospital site to the laboratory for further processing.

#### Mouse breeding and husbandry

All procedures were carried out in accordance with the Institutional Animal Care and Use Committee at the Allen Institute for Brain Science or Vrije Universiteit. Animals (<5 mice per cage) were provided food and water ad libitum and were maintained on a regular 12-h light:dark cycle; rooms were kept at 21.1 °C and 45–70% humidity. Mice were maintained on the C57BL/6J background, and newly received or generated transgenic lines were backcrossed to C57BL/6J. Experimental animals were heterozygous for the recombinase transgenes and the reporter transgenes. For details on transgenic lines, age, or other details see Data S1.

#### Tissue processing

Data were obtained from male and female mice between the ages of postnatal day (P)45 and P70. Mice were anaesthetized with 5% isoflurane and intracardially perfused with 25 or 50 ml of 0–4 °C slicing ACSF. Human or mouse acute brain slices (350 μm) were prepared with a Compresstome VF-300 (Precisionary Instruments) or VT1200S (Leica Biosystems) vibrating blade microtome modified for blockface image acquisition (Mako G125B PoE camera with custom integrated software) before each section to aid in registration to the common reference atlas.

Slices were transferred to a carbogenated (95% O_2_/5% CO_2_) and warmed (34 °C) slicing ACSF for 10 min, then transferred to room temperature holding ACSF of the composition 52 (in mM): 92 NaCl, 2.5 KCl, 1.2 NaH_2_PO_4_, 30 NaHCO_3_, 20 HEPES, 25 D-glucose, 2 thiourea, 5 sodium-L-ascorbate, 3 sodium pyruvate, 2 CaCl_2_.4H_2_O and 2 MgSO_4_.7H_2_O for the remainder of the day until transferred for patch clamp recordings. Before use, the solution was equilibrated with 95% O_2_, 5% CO_2_ and the pH was adjusted to 7.3 using NaOH. Osmolality was verified to be between 295–310mOsm kg^-1^.

#### Patch clamp recording

Slices were continuously perfused (2 mL/min) with fresh, warm (32–34 °C) recording ACSF containing the following (in mM): 126 NaCl, 2.5 KCl, 1.25 NaH_2_PO_4_, 26 NaHCO_3_, 12.5 D-glucose, 2 CaCl_2_.4H_2_O and 2 MgSO_4_.7H_2_O (pH 7.3), continuously bubbled with 95% O_2_ and 5% CO_2_. The bath solution contained blockers of fast glutamatergic (1 mM kynurenic acid) and GABAergic synaptic transmission (0.1 mM picrotoxin). Thick-walled borosilicate glass (Warner Instruments, G150F-3) electrodes were manufactured (Narishige PC-10 or Sutter Instruments P-87) with a resistance of 4–5 MΩ. Before recording, the electrodes were filled with ^~^1.0–2.0 μL of internal solution (110 mM potassium gluconate, 10.0 mM HEPES, 0.2 mM ethylene glycol-bis (2-aminoethylether)-N,N,N′,N′-tetraacetic acid, 4 mM potassium chloride, 0.3 mM guanosine 5′-triphosphate sodium salt hydrate, 10 mM phosphocreatine disodium salt hydrate, 1 mM adenosine 5’-triphosphate magnesium salt, 20 μg mL^-1^ glycogen, 0.5 U μl^-1^ RNAse inhibitor (Takara, 2313A), 0.02 Alexa 594 or 488 and 0.5% biocytin (Sigma B4261), pH 7.3). The pipette was mounted on a Multiclamp 700B amplifier headstage (Molecular Devices) fixed to a micromanipulator (PatchStar, Scientifica or Mini25, Luigs and Neumann).

Electrophysiology signals were recorded using an ITC-18 Data Acquisition Interface (HEKA). Commands were generated, signals were processed and amplifier metadata were acquired using MIES (https://github.com/AllenInstitute/MIES/), written in Igor Pro (Wavemetrics). Data were filtered (Bessel) at 10 kHz and digitized at 50 kHz. Data were reported uncorrected for the measured (Neher 1992) – 14 mV liquid junction potential between the electrode and bath solutions.

Before data collection, all surfaces, equipment and materials were thoroughly cleaned in the following manner: a wipe down with DNA away (Thermo Scientific), RNAse Zap (Sigma-Aldrich) and finally with nuclease-free water.

L1 was identifiable in mouse and human brain slices as the neuron sparse region directly between the pial surface and the neuron dense layer 2/3. Neurons within L1 were targeted for patch clamp recordings.

After formation of a stable seal and break-in, the resting membrane potential of the neuron was recorded (typically within the first minute). A bias current was injected, either manually or automatically using algorithms within the MIES data acquisition package, for the remainder of the experiment to maintain that initial resting membrane potential. Bias currents remained stable for a minimum of 1 s before each stimulus current injection. Upon attaining whole cell current clamp mode, the pipette capacitance was compensated and the bridge was balanced.

The voltage response of each cell was recorded in response to a standardized stimulus paradigm described previously (*26*) that included square pulses, ramps and chirps, with the goal of extracting features that could be compared across cells, rather than tailoring each stimulus to the physiological input of that neuron.

##### Nucleus extraction

Upon completion of electrophysiological examination, the pipette was centered on the soma or placed near the nucleus (if visible). A small amount of negative pressure was applied (^~^−30 mbar) to begin cytosol extraction and attract the nucleus to the tip of pipette. After approximately one minute, the soma had visibly shrunk and/or the nucleus was near the tip of the pipette. While maintaining the negative pressure, the pipette was slowly retracted diagonally in the *x* and *z* direction. Slow, continuous movement was maintained while monitoring pipette seal. Once the pipette seal reached >1 GΩ and the nucleus was visible on the tip of the pipette, the speed was increased to remove the pipette from the slice. The pipette containing internal solution, cytosol and nucleus was removed from pipette holder and contents were expelled into a PCR tube containing the lysis buffer (Takara, 634894).

##### Voltage clamp experiments

For a subset of experiments with a high nucleated patch resistance (>1000 MΩ), we measured macroscopic outward ionic currents in voltage clamp. To reduce capacitive artifacts, the pipette containing the nucleus was raised to the upper portion of the bath. For K+ channels, activation curves were constructed from 1 s depolarizing voltage commands (−50 to +70 mV in 10 mV voltage steps) from a holding potential of −90 mV). Linear leakage and capacitive currents were digitally subtracted by scaling traces at smaller command voltages in which no voltage-dependent current was activated. To isolate K+ currents into their phenomenological components (A-type, D-type, non-inactivating), we exploited the voltage dependent properties of the putative channels underlying each component (*60*). The fast-inactivating component (*I*_K*A*_) was inactivated by a brief step to −20 mV followed by a series of 1 sec voltage steps ranging from −50 to 70 mV in 10 mV increments. *I*_KA_ was obtained by digitally subtracting the resultant current from the total current, post hoc. Step depolarization to 70 mV from a holding potential of −20 mV was used to inactivate all K+ current and revealed a sustained current. Subtracting the sustained current from the current used to isolate I_KA_ revealed a slowly inactivating, D type (I_KD_) current. Peak currents were calculated for each voltage step. Conductance values were calculated based on the recorded membrane potentials and a K+ reversal potential at −100 mV. The surface area of the nucleated patch was calculated to obtain current and conductance densities.

##### Quality control

For an individual sweep to be included in analysis, the following criteria were applied: (1) membrane potential within 2 mV of target potential (initial resting potential of cell); (2) bias (leak) current 0 ± 100 pA; and (3) root mean square noise measurements in a short window (1.5 ms, to gauge high frequency noise) and longer window (500 ms, to measure patch instability) <0.2 mV and 0.5 mV, respectively.

For human electrophysiology in the primary dataset, QC filters were also imposed at the cell level to flag cells with >1 GΩ seal recorded before break-in, initial access resistance <1 or >20 MΩ or >25% of the input resistance. Cell recordings failing these tests were manually examined for recording quality and manually passed or failed. Cells also had to have features successfully extracted for long square pulse sweeps at a minimum to be included in analysis. For mouse VISp cells, slightly stricter automated QC values were imposed at the sweep and the cell level, following the original publication.

#### Transcriptomic data collection

##### cDNA amplification and library construction

We performed all steps of RNA-processing and sequencing as described in our previous human Patch-seq studies (*22, 26, 28, 48*). We used the SMART-Seq v4 Ultra Low Input RNA Kit for Sequencing (Takara, 634894) to reverse transcribe poly(A) RNA and amplify full-length cDNA according to the manufacturer’s instructions. We performed reverse transcription and cDNA amplification for 20 PCR cycles in 0.65 ml tubes, in sets of 88 tubes at a time. At least 1 control 8-strip was used per amplification set, which contained 4 wells without cells and 4 wells with 10 pg control RNA. Control RNA was either Universal Human RNA (UHR) (Takara 636538) or control RNA provided in the SMART-Seq v4 kit. All samples proceeded through Nextera XT DNA Library Preparation (Illumina FC-131-1096) using either Nextera XT Index Kit V2 Sets A-D(FC-131-2001,2002,2003,2004) or custom dual-indexes provided by Integrated DNA Technologies (IDT). Nextera XT DNA Library prep was performed according to manufacturer’s instructions except that the volumes of all reagents including cDNA input were decreased to 0.2× by volume. Each sample was sequenced to approximately 1 million reads.

##### RNA-seq data processing

Fifty-base-pair paired-end reads were aligned to GRCh38.p2 using a RefSeq annotation gff file retrieved from NCBI on 11 December 2015 for human and to GRCm38 (mm10) using a RefSeq annotation gff file retrieved from NCBI on 18 January 2016 for mouse (https://www.ncbi.nlm.nih.gov/genome/annotation_euk/all/). Sequence alignment was performed using STAR v2.5.353 in two pass Mode. PCR duplicates were masked and removed using STAR option bamRemoveDuplicates. Only uniquely aligned reads were used for gene quantification. Gene counts were computed using the R Genomic Alignments package summarizeOverlaps function using IntersectionNotEmpty mode for exonic and intronic regions separately 54. Expression levels were calculated as counts of exonic plus intronic reads. For most analyses, log_2_(counts per million (CPM) + 1)-transformed values were used, or CPM in the case of Seurat or dprime analyses.

#### Anatomical annotations

##### Layer annotation and alignment

To characterize the position of biocytin-labeled cells, a 20× brightfield and fluorescent image of DAPI (4′,6-diamidino-2-phenylindole) stained tissue was captured and analyzed to determine layer position (human and mouse) and region (mouse only). Using the brightfield and DAPI image, soma position and laminar borders were manually drawn for all human neurons and reconstructed mouse cells and were used to calculate depth relative to the pia, white matter, and/or laminar boundaries. Laminar locations were calculated by finding the path connecting pia and white matter that passed through the cell’s soma coordinate, and measuring distance along this path to laminar boundaries, pia and white matter. For mouse cells without reconstructions, pia and white matter boundaries from the CCF were used as references, with layer positions calculated by aligning the relative cortical depth to an average set of layer thicknesses.

For reconstructed neurons, laminar depths were calculated for all segments of the morphology, and these depths were used to create a “layer-aligned” morphology by first rotating the pia-to-WM axis to vertical, then projecting the normalized laminar depth of each segment onto an average cortical layer template.

##### CCF pinning and alignment

Mouse cells were individually manually placed in the appropriate cortical region and layer within the Allen Mouse Common Coordinate Framework (CCF) (*61*) by matching the 20× image of the slice with a “virtual” slice at an appropriate location and orientation within the CCF.

##### Human brain region pinning

Available surgical photodocumentation (MRI or brain model annotation) is used to place the human tissue blocks in approximate 3D space by matching the photodocumentation to a MRI reference brain volume “ICBM 2009b Nonlinear Symmetric” (Fonov et al 2009), with Human CCF overlayed (*62*) within the ITK-SNAP interactive software.

#### Morphological Reconstruction

##### Biocytin histology

A horseradish peroxidase (HRP) enzyme reaction using diaminobenzidine (DAB) as the chromogen was used to visualize the filled cells after electrophysiological recording, and 4,6-diamidino-2-phenylindole (DAPI) stain was used to identify cortical layers as described previously (*28*).

##### Imaging of biocytin-labelled neurons

Mounted sections were imaged as described previously9. In brief, operators captured images on an upright AxioImager Z2 microscope (Zeiss, Germany) equipped with an Axiocam 506 monochrome camera and 0.63× Optivar lens. Two-dimensional tiled overview images were captured with a 20× objective lens (Zeiss Plan-NEOFLUAR 20×/0.5) in bright-field transmission and fluorescence channels. Tiled image stacks of individual cells were acquired at higher resolution in the transmission channel only for the purpose of automated and manual reconstruction. Light was transmitted using an oil-immersion condenser (1.4 NA). High-resolution stacks were captured with a 63× objective lens (Zeiss Plan-Apochromat 63×/1.4 Oil or Zeiss LD LCI Plan-Apochromat 63×/1.2 Imm Corr) at an interval of 0.28 μm (1.4 NA objective) or 0.44 μm (1.2 NA objective) along the *z* axis. Tiled images were stitched in ZEN software and exported as single-plane TIFF files.

##### Morphological reconstruction

Reconstructions of the dendrites and the full axon were generated for a subset of neurons with good quality transcriptomics, electrophysiology and biocytin fill. Reconstructions were generated based on a 3D image stack that was run through a Vaa3D-based image processing and reconstruction pipeline (*63*). For some cells images were used to generate an automated reconstruction of the neuron using TReMAP (*64*). Alternatively, initial reconstructions were created manually using the reconstruction software PyKNOSSOS (https://www.ariadne.ai/) or the citizen neuroscience game Mozak (*65*) (https://www.mozak.science/). Automated or manually-initiated reconstructions were then extensively manually corrected and curated using a range of tools (for example, virtual finger and polyline) in the Mozak extension (Zoran Popovic, Center for Game Science, University of Washington) of Terafly tools (*66, 67*) in Vaa3D. Every attempt was made to generate a completely connected neuronal structure while remaining faithful to image data. If axonal processes could not be traced back to the main structure of the neuron, they were left unconnected.

#### Slice immunohistochemistry

##### Immunohistochemistry and slide imaging

Tissue slices (350 μm-thick) designated for histological profiling were fixed for 2–4 days in 4% paraformaldehyde (PFA) in phosphate-buffered saline (PBS) at 4 °C and transferred to PBS, 0.1% sodium azide for storage at 4 °C. For human samples, these slices were interspersed with patch-seq slices when preparing each tissue block, while for mouse the histology slices were from separate tissue blocks prepared following the same protocol. Slices were then cryoprotected in 30% sucrose, frozen and resectioned at 30 μm (human tissue) and 20 μm (mouse tissue) using a sliding microtome (Leica SM2000R). Sections were stored in PBS with azide at 4 °C in preparation for immunohistochemical staining. Staining for Neu-N (Neuronal nuclei; Millipore, MAB377, 1:2,000) with DAB was applied using the Biocare Intellipath FLX slide staining automated platform (as previously in (*28*)). The images of stained subsections were acquired with 20x objective on Aperio microscope at a resolution of 1 μm to 1 pixel (human slices) and 1 μm to 0.989 (mouse slices). Full immunohistology protocol details available at http://help.brain-map.org/download/attachments/8323525/CellTypes_Morph_Overview.pdf?version=4&modificationDate=1528310097913&api=v2

##### Cytoarchitecture analysis

Images of Neu-N stained sections from human MTG (1 section per donor for 5 donors) and mouse VISp (1 section per mouse for 3 mice) were selected for further analysis. The quantification of cell densities and cell body sizes was performed using custom MATLAB scripts (R2022a, Mathworks). Within each subsection, several regions of interest (ROIs) containing only L1 were selected manually, each ROI a rectangle of 500-700 μm in length and the full extent of L1 in height. The border between L1 and L2 was visually identified as a characteristic sharp increase in cell body size and density. The MATLAB image processing scripts identified the cells by binarizing the image and applying watershed transform. Cell area was extracted after applying the scaling factor (1 μm/pixel for human and 0.989 μm/pixel for mouse). The cell densities were calculated as the number of cells divided by the volume of tissue (the product of selected ROI area and tissue subsection thickness: 30 μm for human tissue and 20 μm for mouse tissue). Statistical tests were performed using Kruskal Wallis test with post hoc comparisons.

#### Transcriptomic data analysis

##### Reference data from dissociated cells and nuclei

Reference transcriptomic data used in this study were obtained from dissociated inhibitory cells (mouse) or nuclei (human) collected from human MTG (*21*) and mouse VISp (*43*), and are publicly accessible at the Allen Brain Map data portal (https://portal.brain-map.org/atlases-and-data/rnaseq). Layer 1 t-types were assessed by proportions in these datasets after first reducing sampling bias by selecting only samples with even dissections across all cortical layers, and in mouse additionally restricting to samples targeted by pan-neuronal or pan-GABAergic mouse lines. L1 t-types were defined as types making up >5% of L1 cells or with >50% of the type found in L1. For human t-types (with relatively unbiased patch-seq sampling), we verified borderline t-types with the more precise layer boundaries in patch-seq data, excluding the VIP LBH type (<1% of L1, <25% in L1) and including the PAX6 TNFAIP8L3 type (>1% of L1, >50% in L1).

##### Discriminant analysis (*d*′)

For measuring distinctness between types in transcriptomic feature space, we followed the cross-validated negative binomial (NB) discriminant analysis from(*23*). For each pair of t-types, a set of cross-validated log-likelihood ratios (LLR) were calculated for each cell, fitting a NB classifier to the training split and measuring LLR for the test split across rounds of 5-fold cross-validation. The classifier was a naive Bayes negative binomial model, with independent negative binomial distributions fit for each subset on each feature (gene) by maximum likelihood, with dispersion parameter set to *r*=1 following the observed statistics of our dataset

This produced a distribution of likelihood ratios for each t-type in the pair, the separation of which was summarized by the *d*′ statistic. For normal distributions this is typically calculated as the separation of means divided by the standard deviation, but we instead used a non-parametric form (equivalent in the normal distribution case): 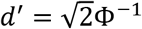 (*AUC*), where Φ is the CDF of the standard normal distribution, and AUC is the area under the receiver operating characteristic curve for the classifier (equivalently, the proportion of pairs selected one from each type for which the LLR of the cluster 1 cell is higher than the cluster 2 cell).

This method was adapted for other modalities (patch-seq transcriptomics, morphology, and electrophysiology) by simply using different classifiers. For electrophysiology and morphology, we used a random forest classifier with scikit-learn default parameters and balanced class weighting. For patch-seq transcriptomics, we modified the naive Bayes negative binomial model to use a zero-inflated negative binomial distribution (statsmodels). Given the high number of free parameters in this model, it was not directly suitable to fitting on the small datasets necessary for the pairwise discriminant analysis. Using ZINB fits to each gene across the full reference dataset, we observed that the parameters π (zero-inflation probability) and ϕ (NB dispersion or shape parameter) both nearly followed a curve depending on μ, the NB distribution mean. We parametrized these curves (ϕ with a spline fit, π with a sigmoid), and used them to constrain ZINB fits for discriminant analysis, essentially assuming that for all genes, zero-inflation parameters follow the same dependence on mean expression. Given these constraints, the maximum likelihood fit for each gene could be implemented simply as a lookup table.

##### Subclasses and cross-species homology

Human L1 transcriptomic subclasses were defined based on the dprime values by grouping all pairs of t-types with dprime<2.2, equivalent to approximately 2% overlap of LLR distributions. Four pairs of t-types were grouped in this way, forming 3 subclasses.

We defined cross-species homology of L1 t-types following a variation of the procedure in (*21*), using coordinates for each cell in a space integrating the mouse and human transcriptomic references (calculated in the original from ScAlign, 30 dimensions). Instead of relying on defining clusters in that integrated space and measuring overlap within those clusters, we directly defined a similarity metric for any two clusters in that space: the ratio of the mean intra-cluster difference to the mean inter-cluster difference. Intra-cluster difference was averaged over all pairs of cells within each cluster, then averaged over the two clusters; inter-cluster difference was averaged directly over all inter-cluster pairs. We summarized this similarity metric for all mouse L1 t-types aligned to human L1 subclasses (Fig 1) and to individual human L1 t-types (Fig S1).

##### Patch-seq data curation and mapping

Patch-seq samples were included in this dataset if they met the following transcriptomic quality criteria: a normalized sum of ‘on’ type marker gene expression (NMS) greater than 0.4 and a normalized sum of non-neuronal contamination markers less than 2.

We mapped Patch-seq samples to reference taxonomies from the reference single cell/nuclei RNA-sequencing datasets introduced above, consisting of a hierarchical dendrogram of cell types, with a subset of cells from the reference identified with each node of the tree and a set of marker genes defined to distinguish types at each split in the tree. The Patch-seq transcriptomes were mapped to the reference taxonomy following the ‘tree mapping’ method (map_dend_membership in the scrattch.hicat package). Briefly, at each branch point of the taxonomy we computed the correlation of the mapped cell’s gene expression with that of the reference cells on each branch, using the markers associated with that branch point (i.e., the genes that best distinguished those groups in the reference), and chose the most correlated branch. The process was repeated until reaching the leaves of the taxonomy (t-types). To determine the confidence of mapping, we applied 100 bootstrapped iterations at each branch point, and in each iteration 70% of the reference cells and 70% of markers were randomly sampled for mapping. The percentage of times a cell was mapped to a given t-type was defined as the mapping probability, and the highest probability t-type was assigned as the mapped cell type.

Only cells mapping to the identified L1 t-types were included in subsequent morpho-electric feature analysis. Neurons from non-L1 t-types present in human L1 were also included in the L1 proportion analysis, and those for which morphological reconstructions were available were included in the supplementary morphology gallery. Some additional quality filters were applied to the mouse VIS cells only, following the procedure in their original publication: excluding cells with poor RNA amplification and “inconsistent” cells as defined by unexpected patterns of mapping probabilities.

##### Joint visualization

We visualized transcriptomic diversity using a nonlinear projection of a transcriptomic space following integration of each species’ patch-seq and reference dissociated cell/nuclei datasets. We first excluded genes potentially related to technical variables: X and Y chromosome genes, mitochondrial genes [Human MitoCarta2.0], and genes most highly expressed in a non-neuronal cell type in the reference dataset. For human patch-seq samples, which had more variable quality of transcriptomic data, we additionally excluded a small set of immune/glial activation-related genes that were shown to introduce non-cell-type-related variability (*68*).

This filtered gene set was loaded and processed by the Seurat pipeline (*30*): expression values were first normalized by the SCTransform model, then the 3000 most variable genes were transformed by CCA and nonlinear warping to integrate the patch-seq and reference datasets (functions FindIntegrationAnchors and IntegrateData). For human patch-seq samples only, the SCTransform normalization additionally reduced effects of contamination by regressing against the normalized contamination marker sum for each cell. The integrated space was then transformed by PCA (30 PCs) followed by UMAP projection (to 2 dimensions) for visualization.

#### Electrophysiology feature analysis

For all electrophysiology stimuli that elicited spiking, action potentials were detected by first identifying locations where the smoothed derivative of the membrane potential (d*V*/d*t*) exceeded 20 mV ms^-1^, then refining on the basis of several criteria including threshold-to-peak voltage, time differences and absolute peak height. For each action potential, threshold, height, width (at half-height), fast after-hyperpolarization (AHP) and interspike trough were calculated (trough and AHP were measured relative to threshold), along with maximal upstroke and downstroke rates d*V*/d*t* and the upstroke/downstroke ratio (that is, ratio of the peak upstroke to peak downstroke).

Following spike detection, summary features were calculated primarily from sweeps with long square pulse current injection: subthreshold properties such as input resistance, sag, and rheobase; spike train properties such as f-I slope, ISI CV, irregularity ratio, and adaptation index; and single spike properties of the first action potential such as upstroke-downstroke ratio, after-hyperpolarization, and width. All relevant features were calculated for both the rheobase sweep and a stimulus ^~^40pA above rheobase. For spike upstroke, downstroke, width, threshold, and inter-spike interval (ISI), ‘adaptation ratio’ features were calculated as a ratio of the spike features between the first and third spike (on the first stimulus to elicit at least 4 spikes). Spiking properties were also calculated for short (3 ms) pulse stimulation and a slowly increasing current ramp stimulus. A subset of cells also had subthreshold frequency response characterized by a logarithmic chirp stimulus (sine wave with exponentially increasing frequency), for which the impedance profile was calculated and characterized by features including the peak frequency and peak ratio. Feature extraction was implemented using the IPFX python package (https://github.com/AllenInstitute/ipfx); custom code used for chirps and some high-level features will be released in a future version of IPFX.

#### Morphology feature analysis

Prior to morphological feature analysis, reconstructed neuronal morphologies were expanded in the dimension perpendicular to the cut surface to correct for shrinkage (*69, 70*) after tissue processing. The amount of shrinkage was calculated by comparing the distance of the soma to the cut surface during recording and after fixation and reconstruction. For mouse cells, a tilt angle correction was also performed based on the estimated difference (via CCF registration) between the slicing angle and the direct pia-white matter direction at the cell’s location (*71*). Features predominantly determined by differences in the *z*-dimension were not analyzed to minimize technical artifacts due to *z*-compression of the slice after processing.

Morphological features were calculated as previously described (*71*). In brief, feature definitions were collected from prior studies (*72, 73*). Features were calculated using the skeleton keys python package (https://github.com/AllenInstitute/skeleton_keys). Features were extracted from neurons aligned in the direction perpendicular to pia and white matter. Laminar axon distribution (bin size of 5 microns) and earth movers distance features require a layer-aligned version of the morphology where node depths are registered to an average interlaminar depth template.

#### Statistical analysis of variability

Unless otherwise specified, statistical analyses were implemented in python using the statsmodels package, and clustering and classification methods implemented using scikit-learn.

##### Variation by subclass and species

To assess the variability of morpho-electric features by subclass within species, we used a one-way ANOVA on ranks (Kruskal-Wallis test) for each feature by subclass Results were reported as fraction of variance explained (ε^2^) and KW test *P*-value. P values were corrected for false discovery rate (FDR, Benjamini–Hochberg procedure) across all features for each data modality. Post-hoc Dunn’s tests were run across all pairs of subclasses (excluding ungrouped t-types), and results FDR-corrected. Analysis of feature relationships with other variables including cell depth, brain region, or donor characteristics were likewise assessed by Mann-Whitney tests for binary variables, KW test for categorical, and Spearman’s or Pearson’s correlations for continuous variables, all FDR-corrected across features by modality.

For cross-species analysis, samples were restricted to only cells present within L1 and belonging to one of the homologous subclasses in each species. Overall cross-species variation was assessed by a Mann-Whitney test for each feature, ranked by effect size *r*, and FDR-corrected by modality. Subclass-dependence of these differences were assessed by a two-way ANOVA on species and subclass (heteroskedasticity-corrected) – where species-subclass interactions were found along with species differences, this was followed by post-hoc tests for species differences within each subclass (MW test).

##### Clustering and classification

Clustering and classification tasks required some preprocessing of electrophysiology data to deal with missing and outlier values – we permitted cells with partially incomplete recordings, for instance, to maximize the usage of available data. Preprocessing included outlier removal, data standardization and imputation, after excluding cells with more than 60% of electrophysiological features missing. Extreme outliers were removed first (LocalOutlierFactor <-20); for standardization, features were centered about the median and scaled by interquartile range (RobustScaler); missing values were imputed as the mean of 5 nearest neighbors (KNNImputer).

Following this preprocessing, electrophysiology and morphology classifiers were trained and tested in a pairwise manner, following the discriminant analysis technique described above, as well as on the full multi-class problem of assigning subclass labels to the full dataset based on electrophysiology. For this problem a multi-class logistic regression classifier was used, with balanced class weights. To assess within-dataset performance, repeated stratified 5-fold cross-validation was used, with classifier predictions on test data aggregated across cross-validation folds to calculate a confusion matrix of performance. Performance was additionally assessed on a held out secondary electrophysiology dataset, as described in the main text. To prevent features with dataset dependence from degrading performance, affected features were excluded based on a one-way ANOVA for the effects of dataset (primary or two secondary datasets collected at different sites). Features with p<0.05 or R^2^>0.05 were excluded.

To demonstrate and visualize discrimination based on small subsets of electrophysiology features, we searched for 1- and 2-dimensional feature subspaces in which each subclass clustered separately from all other cells. A 2-cluster Gaussian mixture model was fit to the data in each subspace, and performance assessed by f1 score (harmonic mean of precision and recall) after identifying the cluster that best matched the subset of interest. Results were shown for the highest ranked subspace for each subclass.

#### MERFISH data collection

Human postmortem frozen brain tissue was embedded in Optimum Cutting Temperature medium (VWR,25608-930) and sectioned on a Leica cryostat at −17 C at 10 um onto Vizgen MERSCOPE coverslips. These sections were then processed for MERSCOPE imaging according to the manufacturer’s instructions. Briefly: sections were allowed to adhere to these coverslips at room temperature for 10 min prior to a 1 min wash in nuclease-free phosphate buffered saline (PBS) and fixation for 15 min in 4% paraformaldehyde in PBS. Fixation was followed by 3×5 minute washes in PBS prior to a 1 min wash in 70% ethanol. Fixed sections were then stored in 70% ethanol at 4C prior to use and for up to one month. Human sections were photobleached using a 150W LED array for 72 h at 4C prior to hybridization then washed in 5 ml Sample Prep Wash Buffer (VIZGEN 20300001) in a 5 cm petri dish. Sections were then incubated in 5 ml Formamide Wash Buffer (VIZGEN 20300002) at 37C for 30 min. Sections were hybridized by placing 50ul of VIZGEN-supplied Gene Panel Mix onto the section, covering with parafilm and incubating at 37 C for 36-48 h in a humidified hybridization oven. Following hybridization, sections were washed twice in 5 ml Formamide Wash Buffer for 30 minutes at 47 C. Sections were then embedded in acrylamide by polymerizing VIZGEN Embedding Premix (VIZGEN 20300004) according to the manufacturer’s instructions. Sections were embedded by inverting sections onto 110 ul of Embedding Premix and 10% Ammonium Persulfate (Sigma A3678) and TEMED (BioRad 161-0800) solution applied to a Gel Slick (Lonza 50640) treated 2×3 glass slide. The coverslips were pressed gently onto the acrylamide solution and allowed to polymerize for 1.5h. Following embedding, sections were cleared for 24-48 h with a mixture of VIZGEN Clearing Solution (VIZGEN 20300003) and Proteinase K (New England Biolabs P8107S) according to the Manufacturer’s instructions. Following clearing, sections were washed twice for 5 min in Sample Prep Wash Buffer (PN 20300001). VIZGEN DAPI and PolyT Stain (PN 20300021) was applied to each section for 15 min followed by a 10 min wash in Formamide Wash Buffer. Formamide Wash Buffer was removed and replaced with Sample Prep Wash Buffer during MERSCOPE set up. 100 ul of RNAse Inhibitor (New England BioLabs M0314L) was added to 250 ul of Imaging Buffer Activator (PN 203000015) and this mixture was added via the cartridge activation port to a pre-thawed and mixed MERSCOPE Imaging cartridge (VIZGEN PN1040004). 15 ml mineral oil (Millipore-Sigma m5904-6X500ML) was added to the activation port and the MERSCOPE fluidics system was primed according to VIZGEN instructions. The flow chamber was assembled with the hybridized and cleared section coverslip according to VIZGEN specifications and the imaging session was initiated after collection of a 10X mosaic DAPI image and selection of the imaging area. For specimens that passed minimum count threshold, imaging was initiated and processing completed according to VIZGEN proprietary protocol. Following image processing and segmentation, cells with fewer than 50 transcripts are eliminated, as well as cells with volumes falling outside a range of 100-300um.

##### Gene panel selection

The 140 gene Human cortical panel was selected using a combination of manual and algorithmic based strategies requiring a reference single cell/nucleus RNA-seq dataset from the same tissue, in this case the human MTG snRNAseq dataset and resulting taxonomy (*21*). First, the reference RNA-seq dataset is filtered to only include genes compatible with mFISH. Retained genes need to be 1) long enough to allow probe design (> 960 base pairs); 2) expressed highly enough to be detected (FPKM >= 10), but not so high as to overcrowd the signal of other genes in a cell (FPKM <500); 3) expressed with low expression in off-target cells (FPKM <50 in non-neuronal cells); and 4) differentially expressed between cell types (top 500 remaining genes by marker score (*74*)). To more evenly sample each cell type, the reference dataset is also filtered to include a maximum of 50 cells per cluster. Second, an initial set of high-confidence marker genes are selected through a combination of literature search and analysis of the reference data.

The main step of gene selection uses a greedy algorithm to iteratively add genes to the initial set. To do this, each cell in the filtered reference dataset is mapped to a cell type by taking the Pearson correlation of its expression levels with each cluster median using the initial gene set of size n, and the cluster corresponding to the maximum value is defined as the “mapped cluster”. The “mapping distance” is then defined as the average cluster distance between the mapped cluster and the originally assigned cluster for each cell. In this case a weighted cluster distance, defined as one minus the Pearson correlation between cluster medians calculated across all filtered genes, is used to penalize cases where cells are mapped to very different types, but an unweighted distance, defined as the fraction of cells that do not map to their assigned cluster, could also be used. This mapping step is repeated for every possible n+1 gene set in the filtered reference dataset, and the set with minimum cluster distance is retained as the new gene set. These steps are repeated using the new get set (of size n+1) until a gene panel of the desired size is attained. Code for reproducing this gene selection strategy is available as part of the mfishtools R library (https://github.com/AllenInstitute/mfishtools).

##### Mapping transcriptomic types

Any genes not matched across both the MERSCOPE gene panel and the mapping taxonomy were filtered from the dataset before starting. From there, cluster means were calculated by dividing the number of cells per cluster by the number of clusters collected. Next, we created a training dataset by finding marker genes for each cluster by calculating the Euclidean distance between all clusters and the mean counts of each gene per cluster. This training dataset was fed into a KNN classifier alongside the MERSCOPEs cell by gene panel to iteratively calculate best possible gene matches per cluster.

---

**Table S1.** L1 subclass characteristics and correspondences.

|  | LAMP5 | MC4R | PAX6 | VIP |
| --- | --- | --- | --- | --- |
| **Human** |  |  |  |  |
| Genes | *PTPRT, PTCHD4, SV2C, LAMP5, DBP* | *SORCS1, ASIC2, TSHZ2, FAM19A1, MC4R, NDNF* | *PAX6, RALYL, MAN1A1, ENOX1, NRXN3* | *KCNQ5, LRRTM4, ASIC2, NRXN3, VIP* |
| Ephys | Early spiking, moderate spike frequency adaptation, no or modest sag, high AP up/down ratio | Highest sag with rapid time constant, irregular spiking in subtype | No/modest sag, high membrane tau, high frequency firing at onset (burst firing or strong adaptation in subtypes) | High sag, regular spiking, fast AHP, low AP up/down ratio |
| Morpho | Dense, horizontally elongated axon often confined to L1. Highly branched dendrites. Axon and dendrite offset | Small, compact, ball-shaped axonal arbor | Dense, horizontally elongated axon with long, sparsely branching dendrite. Axon and dendrite offset | Funnel shaped axonal arbor with simple descending branch |
| Previous types | Neurogliaform | Rosehip |  | VIP |
| **Mouse** |  |  |  |  |
| Genes | *Mpped1, Tox3, Sv2c, Lamp5, Ndnf*<br>*(Npy+/Ndnf+)* | *Sorcs1, Pantr1, Npas1, Npas3, Pcdh7*<br>*(Npy-/Ndnf+)* | *Dpy19L1, Pax6, Tspan12, Ano3, Gm2464*<br>*(Ndnf-/Vip-/Chrna7+)* | *Cpne2, Npas1, Enho, Thsd7a, Vip* |
| Ephys | Late spiking, non-adapting | Early spiking, moderate spike frequency adaptation | Depolarizing subthreshold hump, early spiking, moderate spike frequency adaptation | Early spiking, fast adapting, high input resistance |
| Morpho | Dense, horizontally elongated axon often confined to L1 | Horizontally elongated axon often confined to L1, dendrites with wide horizontal extent |  | Multipolar dendrites, descending axons |
| Previous types | Neurogliaform | Canopy cell (subset) | α7 cell, SBC/SBC-like cell (subset) | VIP, SBC/SBC-like cell (subset) |

**Data S1. (separate file) Donor characteristics**

Cell counts and donor metadata for human and mouse donors of patch-seq samples.

**Data S2. (separate file) Human subclass difference statistics**

Results of statistical testing for effects of subclass on morpho-electric features: Kruskal-Wallis ANOVA on ranks with post-hoc Dunn’s test. P-values FDR-corrected (Benjamini-Hochberg).

**Data S3. (separate file) Metadata effect statistics**

Results of statistical testing for effects of donor metadata on morpho-electric features Kruskal-Wallis ANOVA on ranks with post-hoc Dunn’s test for categorical features (brain region), Mann-Whitney test for binary features (medical condition, sex), Spearman’s correlation for continuous features (age). P-values FDR-corrected (Benjamini-Hochberg).

**Data S4. (separate file) Cross-species difference statistics**

Results of statistical testing for effects of species by subclass on morpho-electric features: two-way ANOVA (Type II) sorted by species effect size, with post-hoc Mann-Whitney test for species differences within subclasses in case of significant species-subclass interaction only. P-values FDR-corrected (Benjamini-Hochberg).

**Fig. S1.**
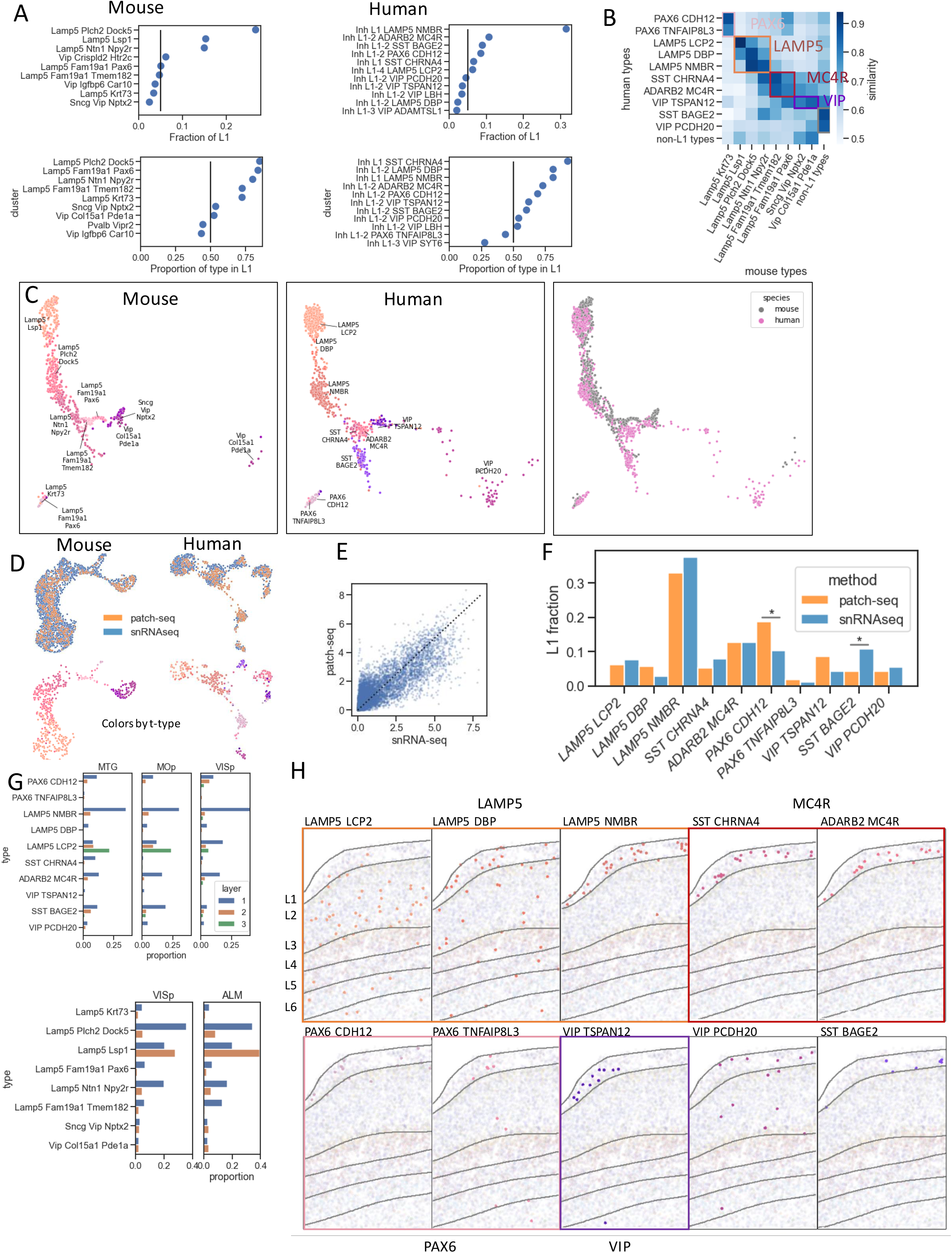
Transcriptomic reference datasets and mapping transcriptomic cell types. A. Criteria for L1 t-types in sn/scRNA-seq reference
B. Cross-species homology of t-types in mouse and human
C. UMAP projection of transcriptomics reference data in aligned transcriptomic space
D. Aligned UMAP projection of patch-seq and snRNA-seq reference, with patch-seq cells labeled by tree mapping classifier.
E. Correlation of marker gene expression between patch-seq and snRNA-seq reference (r=0.84, p<10^-18^).
F. Proportions of t-types in human L1 patch-seq and snRNA-seq reference datasets. Stars indicate p<0.05, FDR-corrected Fisher’s exact test.
G. Proportions of L1 t-types across layer and brain area in human and mouse. Normalized by area, cell counts by layer and t-type divided by total L1 cell count.
H. Spatial distribution of human L1 cell types as revealed by MERFISH. Homology subclasses are denoted by colored boxes. Human t-types with no mouse L1 homology are unboxed.

**Fig. S2.**
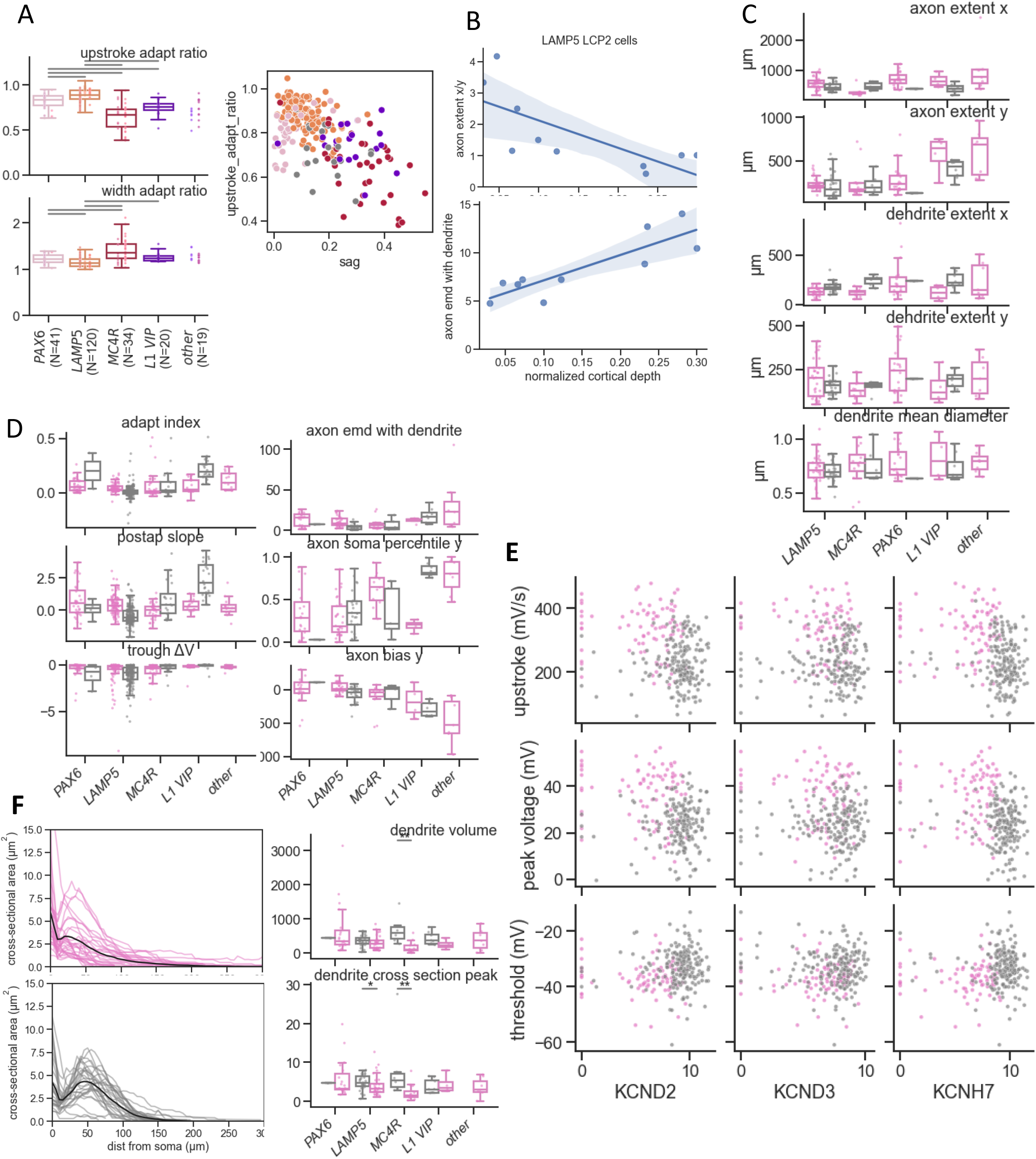
Additional morpho-electric properties of L1 neurons across species. A. Spike adaptation properties of human L1 subclasses, and correlation to sag
B. Variation of axonal arbor shape with cortical depth in the human LAMP5 LCP2 t-type
C. Morphology features relating to overall size showing lack of differences between mouse and human L1 cells
D. Morphology and electrophysiology features with high variation in mouse and not human L1
E. Correlated gene expression and action potential properties differing between mouse and human LAMP5 cells
F. Profiles of total cross-sectional area of dendrites at fixed path length from the soma for individual cells in the LAMP5 subclass (left), showing the effect of increased branching on total volume and peak cross-sectional area (boxplots on right)

**Fig. S3.**
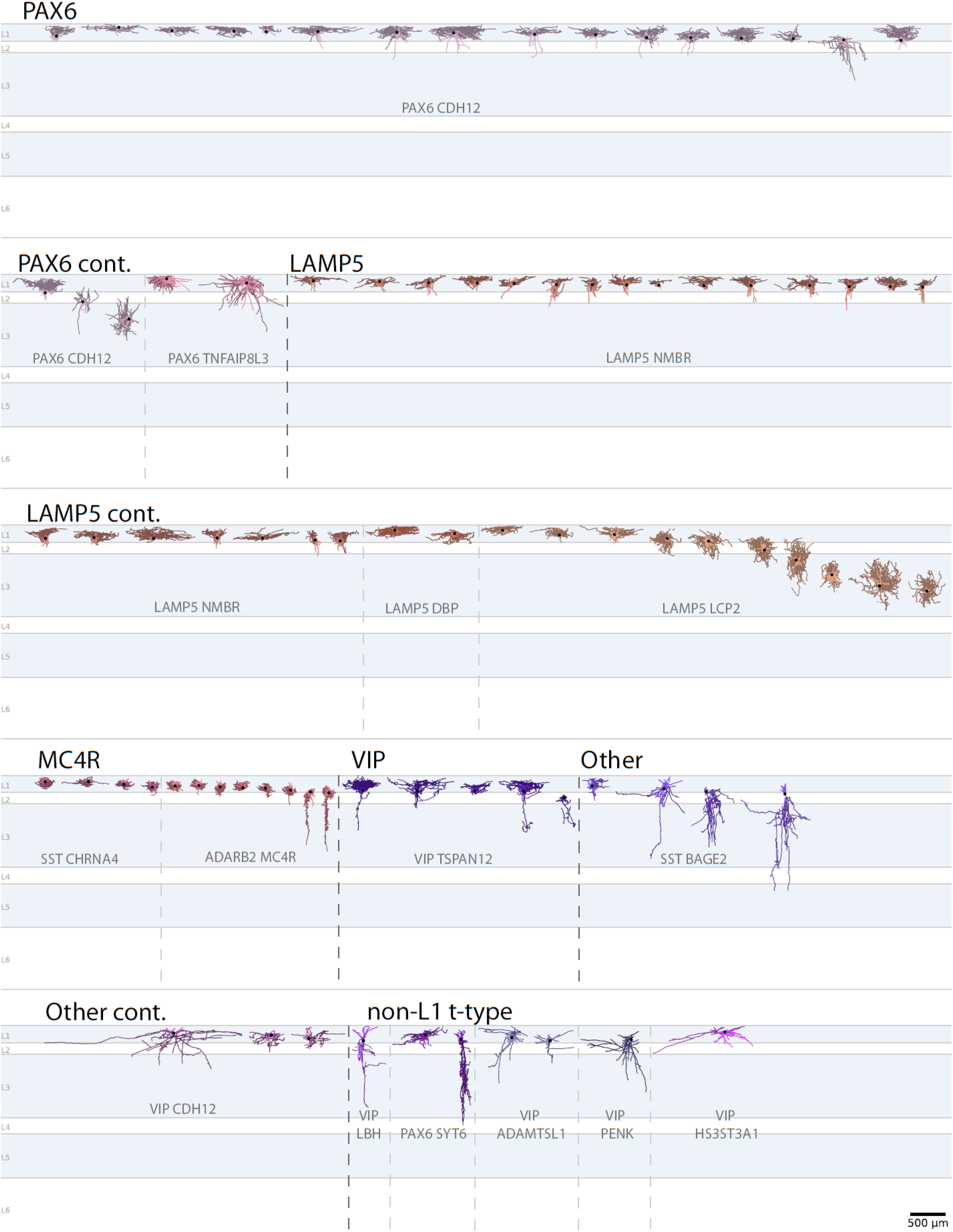
Human morphology gallery. Human dendrite (darker color) and axon (lighter) reconstructions of layer 1 subclasses (black text) and t-types (grey text). Non layer 1 t-types with somas residing in layer 1 also shown.

**Fig. S4.**
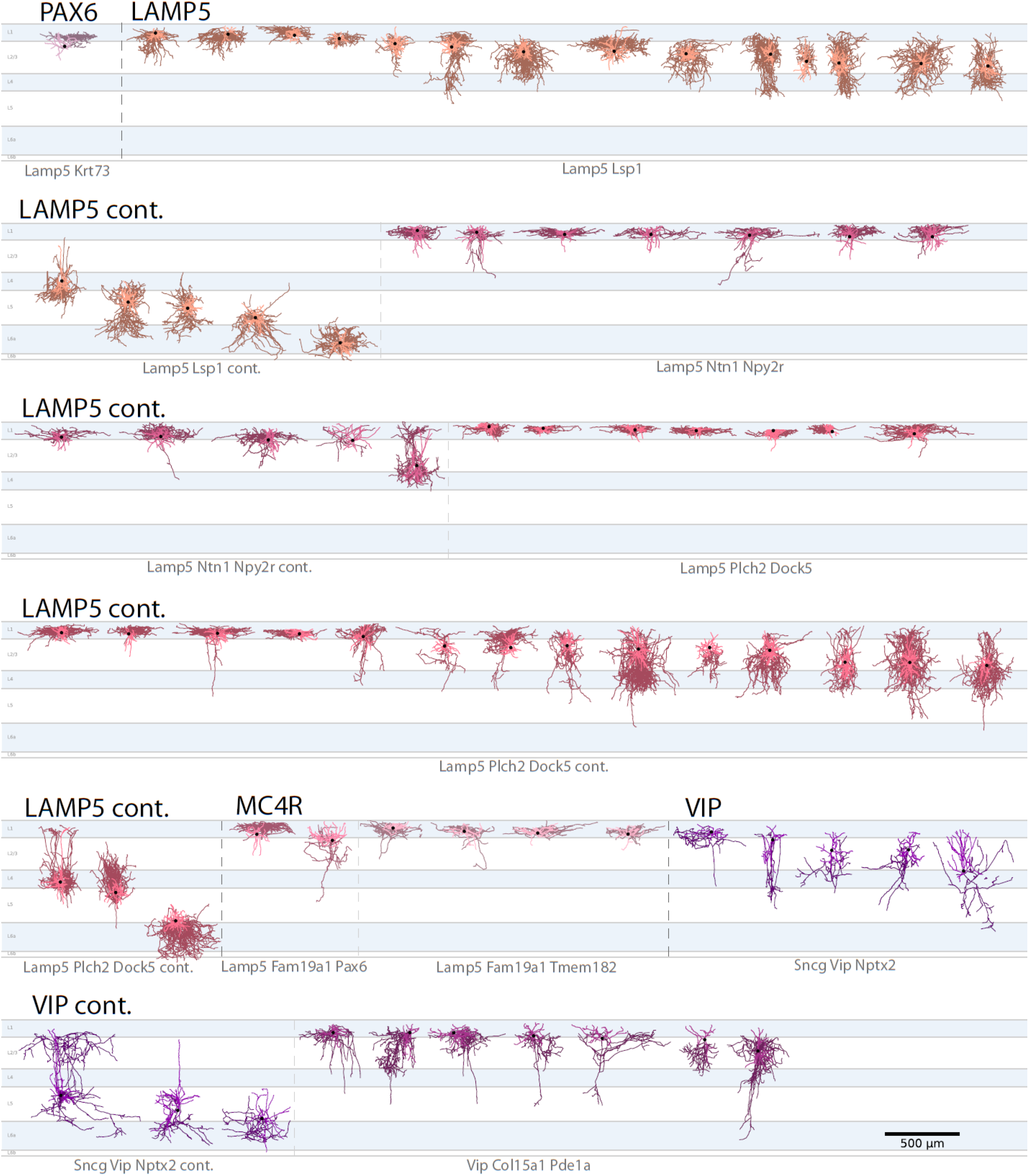
Mouse morphology gallery. Mouse dendrite (darker color) and axon (lighter) reconstructions of layer 1 subclasses (black text) and t-types (grey text).

**Fig. S5.**
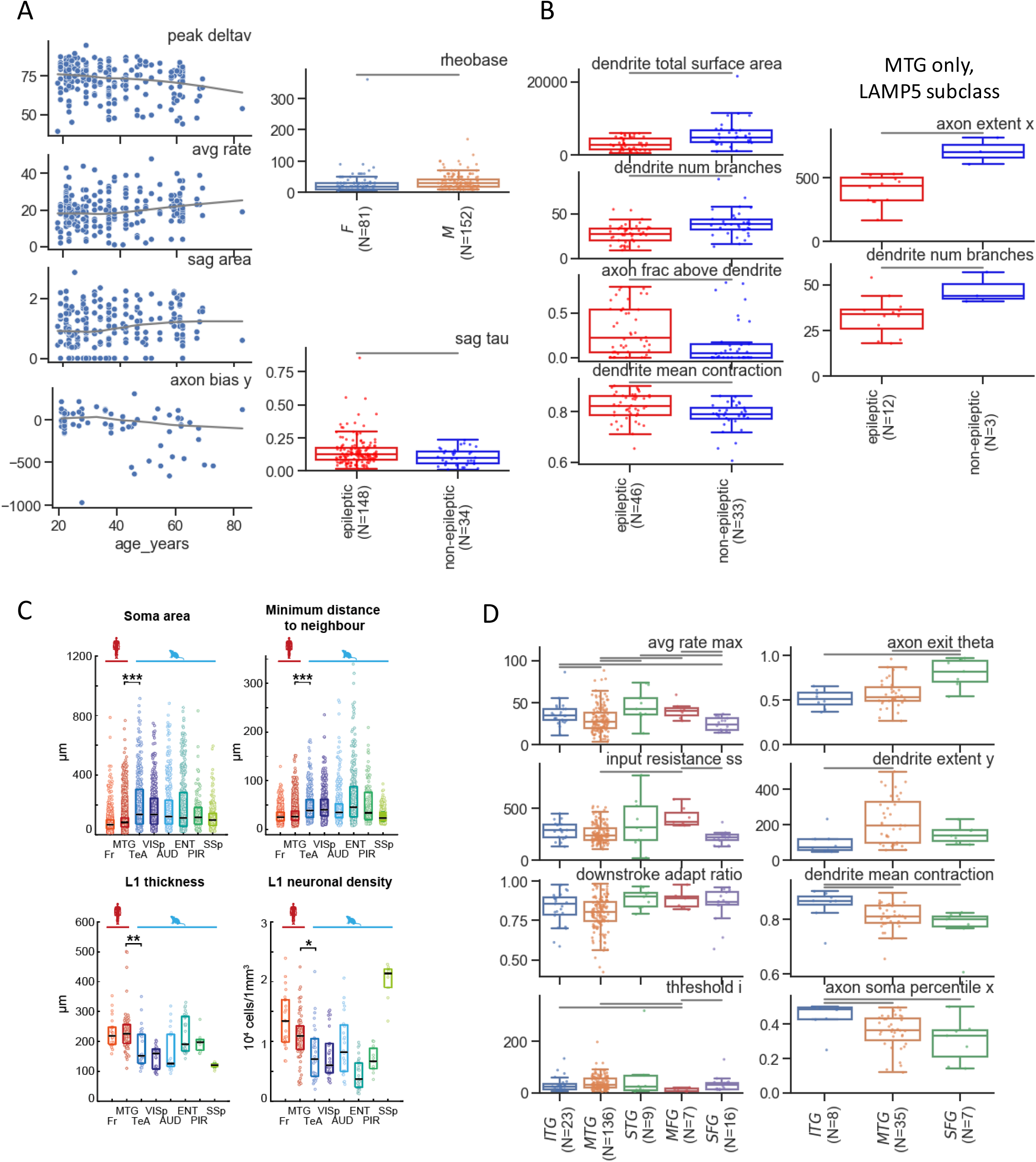
Brain region and donor characteristic effects. A. Minor effects of donor age and sex on L1 neuron morpho-electric features
B. Effects of donor medical condition on morphology features. Although medical conditions are correlated to brain regions sampled, at least one effect remains when testing cells only from the LAMP5 subclass in MTG samples (right)
C. Microstructure quantification across brain regions in mouse and human
D. Electrophysiology and morphology features varying across brain region in human L1 interneuron patch-seq

**Fig. S6.**
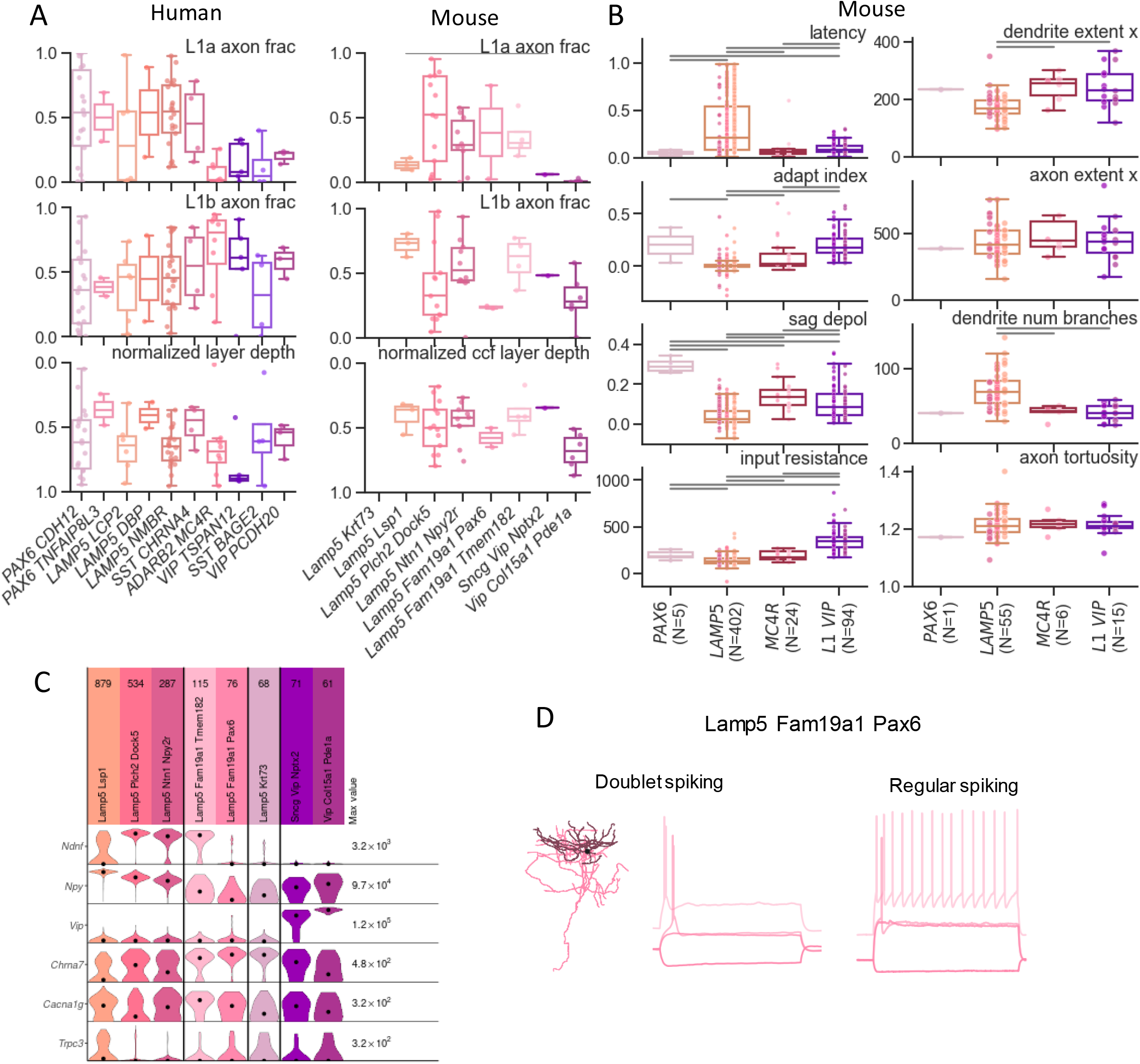
Additional features characterizing MC4R and PAX6 subclasses. A. Sublaminar axon distribution proportions of human (left) and mouse (right) L1 t-types
B. Electrophysiology and morphology features identifying canopy-like cells in mouse L1 MC4R and LAMP5 subclasses
C. Expression of alpha7 type markers and putative bursting-related ion channels in mouse t-types
D. Examples from Lamp5 Fam19a1 Pax6 t-type (MC4R subclass): PAX6-like doublet spiking (left) and regular spiking more characteristic of MC4R subclass (right, no reconstructions available)

**Fig. S7.**
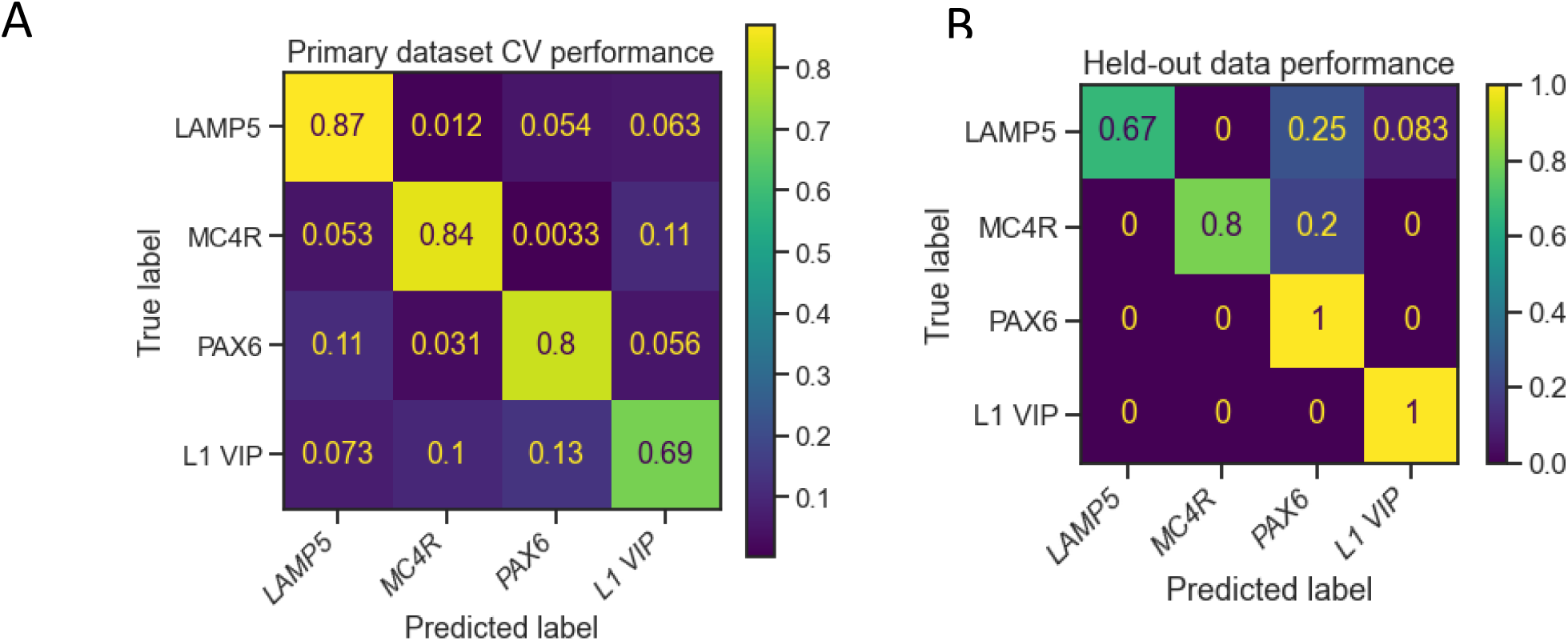
Classifiers can predict human L1 subclasses. Performance assessed using cross-validation (**A**, 82% accuracy) and on held-out test datasets recorded under at different sites (**B**, 81% accuracy)

**Fig. S8.**
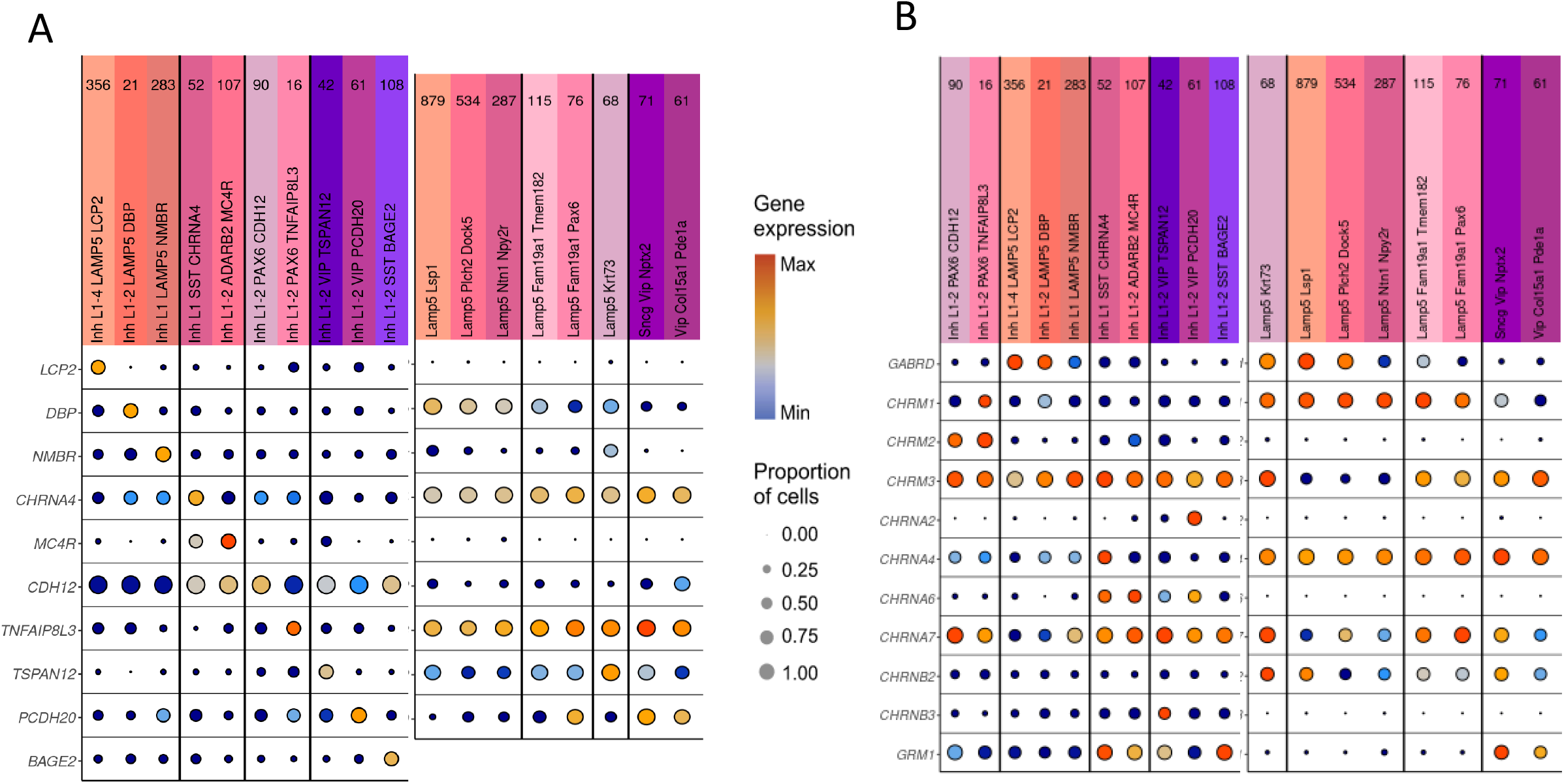
Additional gene expression comparisons for L1 t-types. For both panels, dot size shows the proportion of cells with nonzero expression of a gene, while color shows the median expression in log(CPM+1), normalized by species for each panel.

A. Expression of human t-type marker genes in L1 types in human (left) and mouse (right). Note that the BAGE2 gene has no mouse ortholog.
B. Expression of genes related to neuromodulation and circuit connectivity in L1 types in human (left) and mouse (right)

